# Mixed acid fermentation products from *Lachnospira eligens* counteract myotube atrophy

**DOI:** 10.64898/2026.06.10.729780

**Authors:** Sophie Lecop, Edwige Piron, Audrey M. Neyrinck, Axelle Loriot, Sarah A. Pötgens, Raphaël Helaers, Amandine Jacquet, Pauline Morigny, Eleonora Scorletti, Josh Bilson, Christopher D. Byrne, Patrice D. Cani, Maria Rohm, Lars Vereecke, Thomas C. A. Hitch, Thomas Clavel, Nathalie M. Delzenne, Laure B. Bindels

## Abstract

**Introduction:** Acute myeloid leukemia (AML) is a hematological malignancy associated with muscle wasting. As the relative abundance of *Lachnospira eligens* was reduced in patients with AML compared to healthy individuals and correlated positively with muscle strength, we hypothesized that *L. eligens* positively impacts the muscle through the production of small metabolites reaching the systemic circulation.

**Methods:** *L. eligens* levels were analyzed in two additional independent cohorts. Six *L. eligens* isolates were characterized through whole-genome sequencing to select clinically relevant strains. The composition of their culture supernatant was analyzed by metabolomics. The impact of *L. eligens* supernatant on dexamethasone- and interleukin-6-atrophied murine myotubes was assessed. Bioactive metabolites and their production mechanism were identified using among others bioactivity-guided fractionation. The underlying mechanism was also explored on the host side through myotubes’ transcriptome analysis and metabolic flux analysis. The relevance of bioactive metabolites and their production mechanism was evaluated through clinical data and samples analyses and in a mouse model of leukemia.

**Results:** The levels of *L. eligens* are reduced in independent cohorts of patients with AML and its supernatant counteracts myotube atrophy. This anti-atrophic effect, conserved between strains of the same species, depends on the occurrence of mixed acid fermentation (MAF) in anoxic culture conditions and the presence of its acid end-products acetate, formate and D-lactate. Consistent with those results, blood levels of acetate are decreased and the relative abundance of fecal bacteria capable of performing aerobic respiration is increased in patients with AML. However, bacterial supernatant failed to prevent muscle atrophy and weakness in leukemic mice, likely due to insufficient sustained elevation of acid end-products in the blood.

**Conclusion:** This work reveals the anti-atrophic effect of MAF end-products on myotubes and suggests the importance of considering gut electron acceptor levels (e.g. O_2_) in disorders affecting muscle health.

**Graphical abstract:** Mixed acid fermentation products from *Lachnospira eligens* counteract myotube atrophy.
Our study suggests that gut anaerobiosis is disrupted in treatment-naïve patients with acute myeloid leukemia (AML), leading to decreased circulating acetate levels and a reduced relative abundance of *L. eligens*, which significantly correlated with muscle strength. In line with this framework, *in vitro* experiments demonstrate that the culture supernatant of *L. eligens*, which contains mixed acid fermentation (MAF) end-products such as acetate, effectively counteracts C2C12 myotube atrophy in the presence of pro-atrophying stimuli. Further mechanistic experiments indicate a causal role for MAF end-products in this anti-atrophying effect. Created with BioRender.com. Legend: solid frames: experimental results; dashed frames: hypothetical conclusions derived from results; black solid arrow: established correlation; black dashed arrows: hypothetical causation.

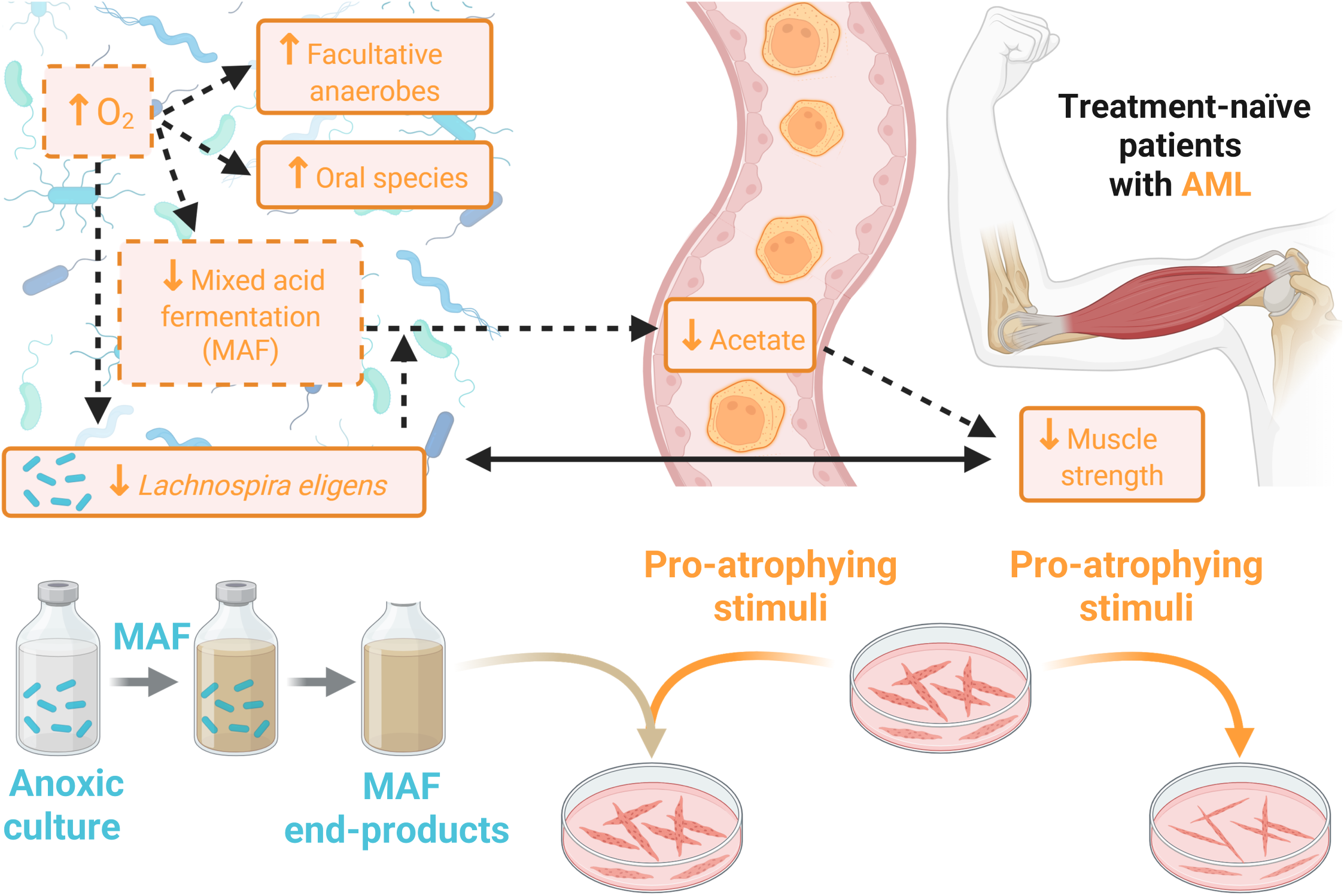

## Introduction

Acute myeloid leukemia (AML) is a hematological malignancy characterized by the uncontrolled proliferation of abnormally differentiated blood myeloid cells that cannot carry out their normal function, leading to anemia and fatigue, hemorrhage, severe infections and in some cases extramedullary disease [1]. Moreover, AML has been associated with low muscle quantity and strength, both of which are induced by muscle wasting and atrophy [2–4]. Muscle atrophy is defined as the biological process of shrinkage of muscle fibers along with changes in fiber types and protein loss, due to the activation of proteolytic systems and the removal of contractile proteins and organelles [5, 6]. Muscle atrophy can occur in localized muscles caused by disuse or denervation but can also occur systemically, the latter being called muscle wasting [7]. Muscle wasting occurs in the elderly, a condition known as primary sarcopenia, as well as in various acute illnesses (burns, sepsis, trauma) and chronic conditions [8]. These chronic conditions include diseases associated with secondary sarcopenia, such as metabolic dysfunction-associated steatotic liver disease (MASLD), as well as cachexia-associated diseases, such as chronic heart failure, chronic obstructive pulmonary disease, chronic kidney disease and cancer [8–10].

Cancer cachexia is defined as a complex metabolic syndrome associated with underlying illness and characterized by loss of muscle, with or without loss of fat mass, that cannot be fully reversed by conventional nutritional support and leads to progressive functional impairment [11, 12]. This multifactorial syndrome occurs in 33.0% of patients with cancer [13] and its prevalence varies depending on cancer type, with pancreatic, gastroesophageal and head and neck cancers having the highest prevalence of cachexia (> 50%) followed by lung, colorectal and hematological cancers (> 40%) [14]. In the context of cancer, low skeletal muscle mass was associated with poorer quality-of-life scores [15] and cachexia or its characteristics have been associated with chemotherapy dose reductions, treatment discontinuation and increased toxicity [16]. The main driving force behind cancer cachexia appears to be systemic inflammation, with a direct effect exerted through several inflammatory cytokines, such as interleukin-6 (IL-6), as well as an indirect effect exerted through the inflammation in the hypothalamus and subsequent increase in circulating glucocorticoids, among others [17, 18]. Both IL-6 and glucocorticoids induce muscle atrophy by increasing MurRF1 and Atrogin-1 expression which leads to the hyperactivation of the ubiquitin-proteasome system [17–20]. Currently, there is no widely approved standard medical treatment for muscle wasting and cachexia. In the case of cancer cachexia, several promising therapies are in late-stage clinical development, and one drug has been approved, anamorelin, but only in Japan [21, 22]. With the aim of developing efficient treatments and since cachexia is a multifactorial syndrome, a great effort has been made in recent years to better understand the different factors influencing it. One of the axes that has attracted interest is the role of the gut microbiome [23–26].

In our previous work, gut dysbiosis was observed in the BaF mouse model of cancer cachexia that mimics acute leukemia [27–29]. In this model, gut dysbiosis was characterized by a decrease in lactobacilli and an increase in *Enterobacteriaceae.* The administration of *Limosilactobacillus reuteri* 100-23 and *Lactobacillus gasseri* 311476 reduced the expression of muscle atrophy markers in the gastrocnemius and in the tibialis, a phenomenon associated with a decrease of inflammatory cytokines [27]. These observations motivated the implementation of the MicroAML study, a multi-centric, prospective, observational clinical study to assess whether the composition and activity of the gut microbiota was affected by AML and whether such changes were associated with metabolic and inflammatory alterations. Fecal, urine and blood samples as well as clinical data, including multiple cachectic parameters, were collected from 30 treatment-naïve antibiotic-free patients with AML, as well as from matched healthy volunteers (1:1). Interestingly, patients with AML showed signs of altered gut function attributed to the leukemia. The bacterium *Lachnospira eligens* (an obligate anaerobic Gram-positive bacterium previously named *Eubacterium eligens* [30, 31]) was found to be reduced by 3-fold in patients with AML and was strongly correlated with muscle strength and citrulline [4], a marker of enterocyte mass and function [32].Several bacterial metabolites have been reported to exert a beneficial effect on muscle physiology, such as short-chain fatty acids (SCFA) and bacterial metabolites arising from the transformation of polyphenols [24, 33]. We therefore hypothesized that *L. eligens* could tackle muscle atrophy through the production of small metabolites that enter systemic circulation and reach the muscle.

To investigate this hypothesis, levels of *L. eligens* were assessed across independent human cohorts. Next, *in vitro* proof-of-concept and mechanistic studies were performed to evaluate the impact of the culture supernatant of clinically relevant rationally selected *L. eligens* strains on myotube atrophy induced by different stimuli relevant to cachexia, namely the synthetic glucocorticoid dexamethasone and IL-6. Finally, the therapeutic potential of these findings was investigated in an *in vivo* experiment and through the analysis of clinical data and samples.

## Materials and Methods

### Gut microbiota analysis

#### Data collection – MicroAML, Wang, Food4Gut and INSYTE (INvestigation of SYnbiotic TreatmEnt in MASLD) cohorts

DNA extraction, metagenomic sequencing data generation and/or 16S rRNA gene sequencing data generation of the MicroAML cohort [4], the Wang cohort [34], the Food4Gut cohort [35] and the INSYTE cohort [36, 37] were described in the original articles. Raw sequences for the four cohorts are present in public repositories: MicroAML cohort: SRA project ID PRJNA813705; Wang cohort: SRA project ID PRJNA746137 (R1 reads, R2 reads sent upon request); Food4Gut cohort: MG-RAST project ID: FOOD4GUThealthy; INSYTE cohort: SRA project ID PRJNA 559052. The same procedures were applied to collect fecal samples, extract DNA, and sequence the 16S rRNA genes for the MicroAML, Food4Gut and INSYTE cohorts. Details of each cohort as well as data processing and analyses are described in the Supplemental Material and Methods.

### Bacterial strains

Six *L. eligens* isolates were gathered from different laboratories and culture collections (Supplemental Table 1). *L. eligens* C15-B4^T^ and *L. eligens* H1-24 were bought from the DSMZ-German Collection of Microorganisms and Cell Cultures GmbH (Germany). *L. eligens* WS1 and *L. eligens* WS2 were kindly provided by the team of Dr. Trevor D. Lawley (Wellcome Sanger Institute, UK) [38]. *L. eligens* I42 was kindly shared by the team of Dr. Petra Louis (Rowett Institute, University of Aberdeen, UK) [39]. *L. eligens* CLA-AA-H260*, Lachnospira hominis* CLA-JM-H89B and *Lachnospira hominis* CLA-JM-H10 were retrieved from the Human Intestinal Bacteria Collection, (HiBC, https://www.hibc.rwth-aachen.de/, Uniklinik RWTH Aaachen University, Germany) [40]. *Escherichia coli* HS was kindly provided by the team of Pr. Mahesh S. Desai (Luxembourg Institute of Health, Luxembourg). The identity of all isolates was validated at the species level by blasting their 16S rRNA gene sequences to the database nucleotide collection (nr/nt) using *Blastn suite* megablast-optimized for highly similar sequences (https://blast.ncbi.nlm.nih.gov/Blast.cgi). To obtain the 16S rRNA gene sequences, universal primers 8F and 1492R were used to amplify 16S rRNA gene, followed by purification and Sanger sequencing performed by Eurofins Genomics Europe Sequencing GmbH (Germany) using primers 27F and 1392R. Primers sequences are reported in Supplemental Table 2.

### Whole-genome sequencing of *Lachnospira eligens* isolates

#### Whole-genome sequencing – data generation

Bacterial whole-genome sequencing was performed on the 6 *L. eligens* isolates gathered from different laboratories and biobanks (Supplemental Table 1). Bacterial pellets were collected by centrifuging (5 minutes, 8 000g, 4°C) 1.5 mL of bacterial culture in 2 mL sterile eppendorfs (Eppendorf, Germany) then stored at -80°C. DNA extraction was performed using the kit DNeasy UltraClean Microbial Kit (QIAGEN, Germany) with the addition of RNase A (QIAGEN, Germany). The pellets were resuspended in 300 µL of PowerBead solution from the kit with the addition of 2 µL of RNase A and the mix was incubated 2 minutes at room temperature. The next steps were followed as stated in the kit. To verify the quality of the extracted DNA, a 1% agarose gel electrophoresis was performed in a buffer solution of Tris Borate EDTA (TEB) buffer 0.5X (Thermo Fisher Scientific, USA) and run at 150 volts. The gel (1% agarose (Eurogentec, Belgium) in TEB 0.5X and 5 µL of SYBR Safe DNA Gel Stain (Thermo Fisher Scientific, USA)) was charged with 15 µL of GeneRuler 1 kb Plus DNA Ladder (Thermo Fisher Scientific, USA) on both sides of the gel as well as 5 µL of sample with 1 µL of Orange DNA Loading Dye (Thermo Fisher Scientific, USA). DNA was then quantified and sent to Eurofins Genomics Europe Sequencing GmbH (Germany) where library preparation was performed followed by whole-genome Sequencing on Illumina NovaSeq X Plus with a 2 x 150 bp paired-end read mode.

### Whole-genome analysis – bioinformatics

Sequence quality assessment as well as *de novo* assembly were performed by Eurofins Genomics Europe Sequencing GmbH (Germany). *Fastp* (version 0.20.0) [41]) was used to trim adapters and low-quality reads (average quality scores < 20) and only reads with a minimum length of 30 bp remained for the downstream analysis. The reads were error corrected and read depth was normalised using *bbnorm* (version 38.76) (https://sourceforge.net/projects/bbmap/). Then *de novo* assembly was performed using *SPAdes* (version 3.15.0) [42] with default parameter settings. Scaffolds and contigs assembly quality was assessed with *QUAST* (version 5.0.2) [43] and visualized with *Icarus* (version 5.0.2) [44].

### Phylogenetic tree

A phylogenetic tree was inferred using *Up-to-date Bacterial Core Genes* (*UBCG*) through the *UBCG* pipeline [45] which is based on the multigene-based phylogenies methods (MBPs), using *prodigal* (version 2.6.3), *hmmer* (version 3.4), *mafft* (version 7.525), *fasttree* (version 2.1.11), *raxml* (version 8.2.13). To visualize the phylogenetic tree of all the *L. eligens* strains, the *Interactive Tree Of Life* (*iTOL*; https://itol.embl.de/) was used.

### Alignment of the scaffolds to the reference genome of the type strain of L. eligens

The scaffolds of each strain were aligned, ordered and oriented to the reference complete genome of the type strain of *L. eligens*, namely *L. eligens* C15-B4^T^ (GCF_000146185.1 ; ASM14618v1 ; RefSeq annotation ; https://www.ncbi.nlm.nih.gov/datasets/genome/GCF_000146185.1/) using *MauveContigMover* [46] with default parameters.

### Assessment of the quality of the aligned genomes

Then the aligned genomes of the six strains were stacked together to be compared using *ProgressiveMauve* [47] with default parameters. Assessment of the quality of the final genomes was performed using *CheckM* (version v1.2.2; lineage specific workflow) and *CheckM2* (version 1.0.2; using *DIAMOND* to annotate genomes) as well as by performing genome annotation with *Bakta* (version 1.9.4; DB type selected full by default). *CheckM* and *CheckM2* enabled to verify that the final genomes were high quality genomes (Supplemental Tables 3 and 4), which are defined by > 90% completeness, < 5% contamination and 10x genome coverage. *Bakta* also enabled to verify that all the final genomes were high quality genomes since they had a detectable 5s rRNA gene sequence, more than 18 unique essential tRNAs genes and detectable 16S and 23S rRNA genes. In addition, *ContEST16S* [48] was used to extract the 16S rRNA gene sequence of the final genomes and blast them, using *Blastn suite*, against the 16S rRNA gene sequences obtained by previously performed full length 16S rRNA Sanger sequencing. The percentage of identity was of 100% for all *L. eligens* strains but *L. eligens* I42 and *L. eligens* WS1 which had a percentage of identity of 99.90% and 99.89% respectively.

### AntiMicrobial Resistance (AMR) genes identification

The final aligned genomes of each of the six strains were queried against six databases [49–54] to identify AMR genes, one database [49] to identify disinfectant resistance genes and one database [55] to identify plasmids which could potentially carry those genes. Different tools were used [49, 50, 55, 56]. The tools and database names, versions and default parameters as well as the date of analysis and the obtained results can be found in the Supplemental Table 5. As one putative AMR gene was identified in *L. eligens* I42 (namely, (Gly)vanA-G), the nucleotide sequence corresponding to (Gly)vanA-G gene in the whole-genome sequence of *L. eligens* I42 was extracted and the reverse strain was translated into an amino acid sequence using *Expasy* (https://web.expasy.org/translate/) to then compare both sequences to the nucleotide and amino acid sequence of (Gly)vanA-G from the *ARG-ANNOT* database using *VectorBuilder* (https://en.vectorbuilder.com). The percentage of identity and coverage of the nucleotide sequences were both 87.34% and for the amino acid sequences were of 84.31% and 87.58% respectively.

#### Overall genome relatedness index (OGRI)

*Orthologous Average Nucleotide Identity Tool* (*OAT*, version 0.93.1) was used to perform OrthoANI [57] on the aligned genomes of the six strains. In addition, digital DNA-DNA hybridization (dDDH) values and confidence intervals were calculated following the recommended settings of the *Genome-to-Genome Distance Calculator 4.0* (*GGDC 4.0*) [58, 59] through the *Type Strain Genome Server* (https://tygs.dsmz.de), an automated high-throughput platform for state-of-the-art genome-based taxonomy [59, 60].

### StrainPhlAn

*StrainPhlAn* (version 4.1) [61] was used to profile *L. eligens* with strain-level resolution in the metagenomics dataset from the MicroAML study and perform phylogenetic comparison of the strains present in the patients and the 6 *L. eligens* isolates. The *CHOCOPhlAnSGB* database (version vJun23_202403) was used. *iTOL* was used to visualize the phylogenetic tree of the *L. eligens* strains present in the same Species Genome Bin (SGB).

### Bacterial growth and supernatant preparation

#### Complex medium preparation protocol

To prepare 1 L of complex medium, 10 g of tryptone (Sigma-Aldrich, USA), 2.5 g of yeast extract (Gibco, USA), 4 g of NaHCO_3_ (Sigma-Aldrich, USA), 2 g of (D)+Glucose (Sigma-Aldrich, USA), 2.10529 g of (D)+Maltose.H_2_O (TCI Europe, Belgium), 2 g of (D)+Cellobiose (Thermo Scientific, USA) and 1.301 g of L-cystine dihydrochloride (Sigma-Aldrich, USA) were mixed in 500 mL of milli-Q water. Then, 150 mL of mineral solution 1, 150 mL of mineral solution 2, 10 mL of hemin solution and 1 mL of resazurin solution were added and mixed. Finally, 189 mL of milli-Q water were added, and the pH was adjusted to 7.45 with a NaOH (Supelco, USA) 10N solution. Then, the medium was heated and mixed (without reaching boiling point) to dissolve all the components and, more importantly, to transfer O_2_ from the liquid phase to the gaseous phase. Finally, 100 mL of medium was transferred into 115 mL vials which were then closed with butyl rubber septa, secured with aluminum caps and flushed with N_2_ gas for 15 minutes at 1.5 bar so that the O_2_ present in the headspace was replaced by N_2_ gas. To sterilize the medium, medium containing vials were autoclaved 15 minutes at 121°C.

To prepare 1 L of mineral solution 1, 3 g of K_2_HPO_4_ (Supelco, USA) were dissolved in 1 L of milli-Q water. To prepare 1 L of mineral solution 2, 3 g of KH_2_PO_4_ (Supelco, USA), 6 g of (NH_4_)_2_SO_4_ (Sigma-Aldrich, USA), 6 g of NaCl (Thermo Scientific, USA), 0.6 g of MgSO_4_ (Sigma-Aldrich, USA) and 0.6 g of CaCl_2_ (dry) (Supelco, USA) were dissolved in 1 L of milli-Q water. To prepare 100 mL of hemin solution, 0.28 g of KOH (EMSURE, Germany) were dissolved in 1,25 mL of milli-Q water and 23.75 mL of ethyl alcohol pure (Sigma-Aldrich, USA). Then, 0.1 g of hemin (Sigma-Aldrich, USA) was added followed by 75 mL of milli-Q water. To prepare 100 mL of resazurin solution, 0.1096 g of resazurin sodium salt (Sigma, USA) was dissolved in 100 mL of milli-Q water. Resazurin was used as a redox indicator to confirm the absence of oxygen in anoxic environments.

### Defined medium preparation protocol

To prepare 1 L of defined medium, 2 g of (NH_4_)_2_SO_4_, 13.6 g of KH_2_PO_4_, 2 g of (D)+Glucose and 1.301 g of L-cystine dihydrochloride were mixed in 500 mL of milli-Q water. Then, 10 mL of 20% glycerol (Sigma-Aldrich, USA) solution, 1 mL of 1M MgSO_4_ solution, 1 mL of 0.5g/L FeSO_4_.7H_2_O (Sigma-Aldrich, USA) solution and 1 mL of resazurin solution were added. Finally, 487 mL of milli-Q water were added, and the pH was adjusted to 7 with an NaOH 10N solution. Then, the medium was heated and mixed (without reaching boiling point). Finally, to obtain anoxic medium, 100 mL of medium was transferred into 115 mL vials which were then closed with butyl rubber septa, secured with aluminum caps and flushed with N_2_ gas for 15 minutes at 1.5 bar so that the O_2_ present in the headspace was replaced by N_2_ gas. To obtain oxic medium, 100 mL of medium was transferred into 500 mL bottles which were not flushed and, when culturing bacteria, were shaken to enhance O_2_ transfer from the headspace to the liquid phase. To sterilize the medium, medium containing vials and bottles were autoclaved 15 minutes at 121°C.

### Lysogeny Broth (LB) preparation protocol

To prepare 1 L of LB medium, 10 g of peptone from casein (Sigma-Aldrich, USA), 5 g of yeast extract and 10 g of NaCl were mixed in 1 L of milli-Q water then autoclaved 15 minutes at 121°C.

### Bacterial stocks preparation protocol

Bacterial glycerol stocks of all *L. eligens* strains, *L. hominis* CLA-JM-H89B and *L. hominis* CLA-JM-H10 were stored at -80°C in small vials. They were obtained by mixing 3 mL of bacterial culture and 2 mL of sterilized glycerol solution in small vials that were previously sterilized and flushed with N_2_ gas. To prepare the glycerol solution, 200 mL of glycerol (Sigma-Aldrich, USA), 200 mL of milliQ water, 0.6 mL of resazurin solution and 0.0230g of L-cystine dihydrochloride (Sigma-Aldrich, USA) were mixed and autoclaved 15 minutes at 121°C.

Bacterial glycerol stocks for *E. coli* HS were stored at -80°C in cryotubes. They were obtained by centrifuging 1 mL of bacterial culture (10 minutes, 5 000g, room temperature), and the pellet was suspended in 300 µL of a sterilized 15% glycerol (Sigma-Aldrich, USA) - 85% PBS (Gibco, USA) solution.

### Bacterial culture

All bacteria were cultivated by first performing a pre-culture that was then used to inoculate new fresh medium and obtain the bacterial culture. All experiments shown were performed using the bacterial culture and not the pre-culture.

Bacterial culture of *L. eligens* strains, *L. hominis* CLA-JM-H89B and *L. hominis* CLA-JM-H10 was performed by first inoculating the content of 5 mL of bacterial stocks in a vial with 100 mL of anoxic culture medium. Following 48 hours, 5 mL of bacterial pre-culture were inoculated in a vial with 100 mL of anoxic culture medium.

Bacterial culture of *E. coli HS* was performed by first inoculating it in 15 mL of aerobic LB medium. Following 24 hours, 5 mL of bacterial pre-culture were inoculated in a vial with 100 mL of anoxic culture medium.

### Optical density measurements

Optical density (OD) was measured at 600 nm in the spectrophotometer SpectraMax i3X (Molecular devices, USA). OD was measured in technical triplicates using 250 µL of bacterial culture in 96-well plates. The shown ODs were obtained by subtracting the OD of the non-conditioned medium (NCM) from the OD of bacterial culture.

### Bacterial culture supernatant and NCM collection

Bacterial culture supernatant and NCM were collected by centrifuging (10 minutes, 5 000g, 4°C) bacterial culture and NCM twice, followed by a filtration at 0.22 microns using Millex-GP Filter Units (Millipore, USA) for sterilization purposes. They were then stored at -80°C.

### Bacterial culture supernatant and NMC treatments

Filtration of the supernatants and NMC was performed using 10 kDa (Amicon Ultra Centrifugal Filter Unit 2 mL, Millipore, Belgium), 3 kDa (Amicon Ultra Centrifugal Filter 0.5 mL, Millipore, Belgium) and 1 kDa (Filter Spin Centrifugal Microsep Advance with Omega membrane, Cytiva, Belgium) molecular-weight-cutoff filters by centrifugation (30 minutes, 4°C) at 4 000g, 14 000g and 5 000g respectively. Filters were pre-washed 2 times with milli-Q water.

To denature proteins, 1 mL of the supernatants and NCM were heated at 95°C during 5 minutes in 2 mL closed safe-lock eppendorfs (Eppendorf, Germany).

To precipitate proteins and eliminate them from the samples, the supernatants and NCM were placed in acetone-compatible tubes and four times the volume of ice-cold acetone (-20°C) was added. Following incubation (60 minutes, -20°C), the samples were centrifuged (10 minutes, 14000g, 25°C). The supernatant was collected and acetone was evaporated until the initial volume of the samples was reached (2 hours, 200bar, 37°C followed by 30 minutes, 120bar, 37°C) using a RapidVap (LABCONCO, Belgium).

To eliminate the acids, acetate, formate and lactate, from the supernatant of *L. eligens* H260 and *E. coli* HS as well as from NCM filtered at 1 kDa, supernatants were run through the acid-binding columns HyperSep SAX Cartridges (Thermo Scientific, USA). To do so, a rack (Agilent Technologies, USA) equipped with a manometer (30 inHg) (Ashcroft, USA) and a vacuum pump (23 inHg in capacity) (VWR, USA) were used. The cartridges were conditioned with 1.5 mL of ethanol (ethyl alcohol pure, Sigma-Aldrich, USA) followed by 3 mL of milli-Q water twice and finally 1 mL of sample. The samples coming out of the cartridges were stored at -80°C and then used for further experiments.

Acetate (Sodium formate, Sigma-Aldrich, Belgium), formate (Sodium formate, Sigma-Aldrich, Belgium) and D-lactate (Sodium D-lactate, Sigma-Aldrich, Belgium) were dissolved and diluted in NCM to obtain a final concentration in NCM of 7.47 mM, 10.51 mM and 2.55 mM respectively. Those concentrations correspond to the mean of the concentration measured in the supernatants of the cultured *L. eligens* strains by ^1^H-NMR metabolomics.

### Cell culture & myotube analyses

C2C12 murine myoblasts were cultured in growth medium containing high glucose Dulbecco’s Modified Eagle Medium (DMEM) high glucose (Gibco, USA) supplemented with 7.96% Fetal Bovine Serum (Biowest, France), 0.88% MEM Non-Essential Amino Acids Solution (Gibco, USA), 88.5 U/ml penicillin-streptomycin (Gibco, USA), and 3.5 mM L-glutamine (Gibco, USA) at 37°C with 5% CO_2_. Once the C2C12 cells reached 80-90% confluence, growth medium was replaced with differentiation medium containing DMEM high glucose (Gibco, USA) supplemented with 1.88 % Horse Serum (Gibco, USA), 0.94 % MEM Non-Essential Amino Acids Solution (Gibco, USA), 100 U/ml penicillin-streptomycin (Gibco, USA), and 4mM L-glutamine (Gibco, USA). After four days of differentiation, cells were treated for 48h with dexamethasone (Sigma-Aldrich, USA) at 1 µM or mouse interleukin-6 recombinant protein (IL-6) (PeproTech, Belgium) at 200 ng/mL and/or NCM collected in our laboratory and/or bacterial culture supernatant collected in our laboratory. For each experiment, the pH of the different C2C12 culture media was verified to ensure that the addition of the various treatment components did not result in pH differences between experimental groups.

For myotube diameter quantification, images were captured using a phase contrast microscopy (EVOS XL Core or EVOS M3000 Imaging System, Thermo Fisher Scientific, Belgium). The myotube diameter was quantified blindly with the image processing software ImageJ (U.S. National Institutes of Health, USA) by a researcher not involved in data analysis. Four pictures were taken of each culture dish . For each picture, 10 myotubes were randomly selected and five measurements for each myotube were performed. Myotubes were defined by the presence of a minimum of 5 nuclei. The transcriptome of C2C12 myotubes was analyzed by RNA sequencing and qPCR, as described in the Supplemental Material and Methods. The experiments were performed in technical triplicates and, when indicated in the legends, also in biological triplicates (three independent experiments).

The Pro-B lymphocyte (BaF3) cell line transfected with Bcr-Abl was a gift from Dr. K. Bhalla (MCG Cancer Center, Medical College of Georgia, Augusta, GA, USA). The BaF cells were maintained as previously described [27].

### Mouse experiments

Forty-eight female Balb/c mice (7 weeks old, Charles River Laboratories, Italy) were kept in specific pathogen-free conditions and housed 2 mice per cage in individually ventilated cages with a 12 h light/dark cycle and fed an irradiated chow diet (AO4-10, SAFE, Tecnilab-BMI, The Netherlands). The experiment was composed of 4 groups of 12 female mice. One cage of the group BaF_Ecoli_OX was excluded from the experiment due to loss following tail vein injection. Following one week of acclimatation and the day before the injection of BaF cells, grip strength was measured using a Grip strength test apparatus (Bioseb, France). The mean of two grip strength measurements per mice (with a 2-minute delay) was considered. At day 0, the first group of mice (CT_NCM_AN) were injected in the tail vein with 100 µL of a sterile solution of NaCl 0.9% and the three other groups (BaF_NCM_AN, BaF_Ecoli_AN and BaF_Ecoli_OX) with 10^6^ Bcr-Abl-expressing BaF3 cells in 100 µL of a sterile solution of NaCl 0.9%. The next 14 days, the groups CT_NCM_AN and BaF_NCM_AN were orally administered with 200 µL of anoxic NCM and the groups BaF_Ecoli_AN and BaF_Ecoli_OX were administered with 200 µL of the supernatant of *E. coli* HS cultured in anoxic or oxic conditions respectively. The administered solutions were concentrated using a CentriVap centrifugal concentrator with a cold trap (LABCONCO, Belgium) and then filtered at 1 kDa. During those 14 days, body weight, food intake and animal well-being were monitored daily. On day 14, grip strength was measured, treatments were orally administered, and animals were then humanely euthanized under deep isoflurane anesthesia (Abbot, USA). Systemic blood was collected from the cava vein in EDTA tubes and centrifuged (13 000 g, 3 min, 4°C). Tissues were weighed and frozen in liquid nitrogen. All samples were stored at −80 °C. Tissue mRNA analysis was performed by qPCR and is described in the Supplemental Material and Methods. All the experiments were performed following the ARRIVE guidelines 2.0 and were approved by and performed in accordance with the guidelines of the local ethics committee from the UCLouvain, Belgium.

### Statistical analysis

Statistical analyses were performed using *GraphPad Prism* (version 8.0.1) (GraphPad Software, USA). P < 0.05 was considered statistically significant. Normality of the results of the *in vivo* experiment and human cohorts was assessed using the Shapiro-Wilk normality test.

When comparing groups two by two and if data distribution was normal, a Student t test was performed whereas if data distribution was not normal, a Mann-Whitney U test was performed.

When comparing more than two groups and if data distribution was normal, the Brown-Forsythe test was performed to evaluate the equality of standard-deviations (SDs) between groups. In case of equal SDs, one-way ANOVA was performed followed by a Dunnett’s pairwise comparison post-test to compare to the positive control group treated with dexamethasone and NCM or a Tukey’s pairwise comparison post-test to compare all the groups. In case of non-equal SDs, a Welch’s ANOVA test was performed followed by a Tamhane T2’s post-test to compare all the groups.

When comparing more than two groups, if data distribution was normal and the data was paired (biological triplicates or food intake and body weight evolution), a repeated measures ANOVA was performed. In case of missing values, a mixed-effects model with the Geisser-Greenhouse correction was built. Those tests were followed by a Dunnett’s or Tukey’s pairwise comparison post-test as indicated in the legends.

When comparing more than two groups and if data distribution was not normal, a Kruskal-Wallis test was performed followed by a Dunnett’s or Tukey’s pairwise comparison post-test as indicated in the legends.

## Results

### The relative abundance of *L. eligens* is decreased in treatment-naive patients with AML

The relative abundance of *L. eligens* was decreased in Belgian patients with AML (MicroAML study), at the time of diagnosis prior to any intervention, compared to paired healthy individuals (Figure 1A). The relative abundance of *L. eligens* was also reduced in an independent cohort of Chinese patients with AML (Wang study), at the time of diagnosis prior to any intervention, compared to healthy individuals (Figure 1A). To validate the relative abundance of *L. eligens* within the healthy Belgian population, we compared it against a second cohort (Food4Gut study), which was not significantly different from the healthy patients in the MicroAML study (Figure 1A).

**Figure 1.**
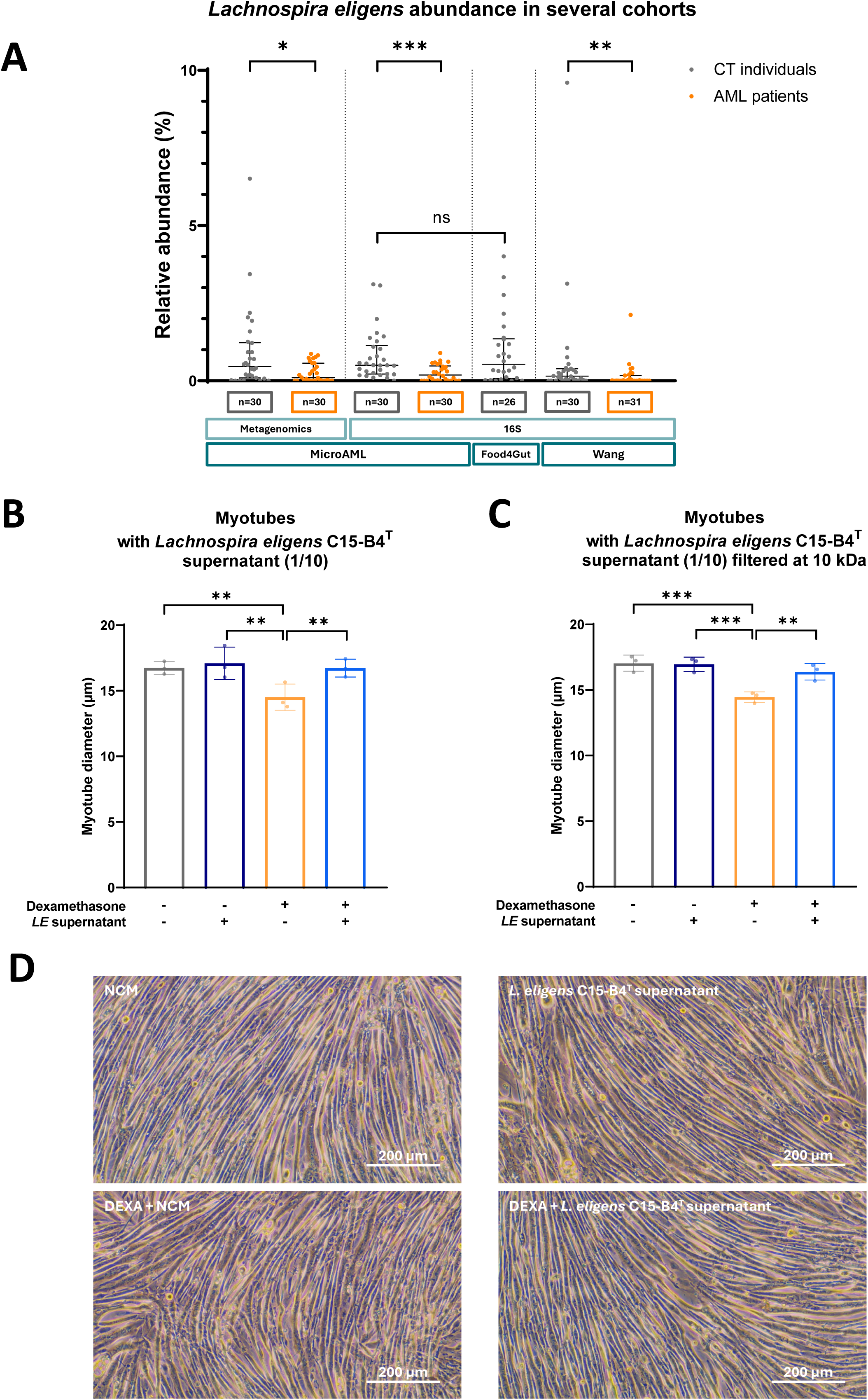
The relative abundance of *Lachnospira eligens* is decreased in patients with AML and the supernatant of *L. eligens* C15-B4^T^ counteracts dexamethasone-induced myotube atrophy. (A) *L. eligens* abundance in treatment-naive AML patients (AML in orange) and healthy individuals (CT in grey) in three independent cohorts : the MicroAML cohort (Pötgens *et al., Haematologica* 2024), the Wang cohort (Wang *et al., Nat Commun* 2022) and the Food4Gut cohort (Hiel, Bindels, Pachikian, Kalala *et al., ACJN* 2019). Mann-Whitney U tests, **p<0.01, ***p<0.001. (B) Comparison of myotube diameter between myotubes incubated with or without 1 µM of dexamethasone and with or without 1/10 diluted *L. eligens* C15-B4^T^ supernatant. Data are presented as mean ± SD, representative of 3 independent experiments performed in triplicates (N = 3, n = 3). Repeated Measures One-Way ANOVA (p=0.0017) followed by Tukey’s multiple comparisons test, **p<0.01. (C) Comparison of myotube diameter between myotubes incubated with or without dexamethasone and with or without 1/10 diluted *L. eligens* C15-B4^T^ supernatant filtered at 10 kDa. Data are presented as mean ± SD, representative of 3 independent experiments performed in triplicates (N = 3, n = 3). Repeated Measures One-Way ANOVA (p=0.0005) followed by Tukey’s multiple comparisons test, **p<0.01, ***p<0.001. (D) Phase contrast microscopy pictures representative of each group of the experiment presented in Figure 1B. Scale bar = 200 µm. Abbreviations : *LE* (*Lachnospira eligens*), NCM (non-conditioned medium), DEXA (Dexamethasone).

### The supernatant of the type strain of *L. eligens* counteracts dexamethasone-induced myotube atrophy

To test the hypothesis that *L. eligens* impacts muscle atrophy through the production of small metabolites, the type strain of *L. eligens*, namely *L. eligens* C15-B4^T^, was cultured in an anoxic complex medium (Supplemental Figure 1A) and the collection of its supernatant was optimized to achieve the maximal concentration of metabolites. To do so, time-series analysis over 78 hours was performed to study the metabolomic profile of culture medium inoculated or not with *L. eligens* C15-B4^T^ using ^1^H-NMR metabolomics (Supplemental Figure 1B). The PCA on metabolomic profile over time showed that the composition of the NCM did not change over time whereas the composition of the supernatant of *L. eligens* C15-B4^T^ varied with time but remained stable from 33 hours on (Supplemental Figure 1B), time at which bacterial culture was in stationary phase (Supplemental Figure 1A). Given those results, the timing for the collection of the bacterial supernatants for further experiments was set to 48h from inoculation.

Next, proof-of-concept *in vitro* experiments were performed to evaluate the effect of *L. eligens* C15-B4^T^ supernatant on C2C12 myotubes treated or not during 48 hours with 1 µM of dexamethasone. Forty-eight hours of incubation with *L. eligens* C15-B4^T^ supernatant had no significant effect on myotube diameter compared to the incubation with non-conditioned medium (NCM) (Figure 1B and 1D), which suggests the absence of a hypertrophic effect of the supernatant of *L. eligens* C15-B4^T^ in healthy conditions. In the presence of dexamethasone, myotube diameter was significantly decreased compared to the incubation with NCM alone, which was counteracted by the supernatant of *L. eligens* C15-B4^T^ (Figure 1B and 1D). To ensure that lipoteichoic acid (∼20 kDa), a major cell wall component of Gram-positive bacteria, did not interfere with the results of the experiment, we performed the same experiment with *L. eligens* C15-B4^T^ supernatant filtered at 10 kDa (Figure 1C). The same results were observed, suggesting that lipoteichoic acid did not modify the anti-atrophic effect of *L. eligens* C15-B4^T^ supernatant.

### The anti-atrophic effect of *L. eligens* is conserved between strains

To study the strain level variation in the anti-atrophic effect of *L. eligens*, 5 additional commensal *L. eligens* isolates available in different laboratories and culture collections were gathered (Figure 2A, Supplemental Table 1). To characterize these 5 isolates alongside the type strain, growth curves were determined (Figure 2B) and their genomes were analyzed. The genomes were screened against 6 antimicrobial resistance (AMR) genes databases, leading to the identification of one putative AMR gene against vancomycin in *L. eligens* I42 (Supplemental Figure 2A). To evaluate if the isolates represented different strains, overall genome relatedness indices were calculated, namely the Orthologous Average Nucleotide Identity (OATANI) (Figure 2C) as well as digital DNA-DNA hybridization (dDDH) (Supplemental Figure 2B). The indices showed a 100% identity between the genomes of *L. eligens* H1-24 and WS2 meaning they were genetically the same strain. In addition, the comparison of the genome of *L. eligens* C15-B4^T^ and the other 5 isolates showed an OATANI lower than 95% and a dDDH lower than 70%, suggesting that these 5 isolates and *L. eligens* C15-B4^T^ represent two different species. Those observations were also made when performing a genome-based phylogenetic tree inference (Supplemental Figure 2C). Then, the genomes were used as references to evaluate whether the 6 isolates were representative of the strains present in the microbiota of the participants from the MicroAML study using *StrainPhlAn*. *L. eligens* C15-B4^T^ belonged to a different Species Genome Bin (SGB) than the other 5 isolates and this SGB was less prevalent in the MicroAML cohort. On the contrary, the SGB to which the other 5 isolates belonged was present in most of the participants (Supplemental Figure 2D). A tree representation of the latter showed that the other 5 isolates were representative of the *L. eligens* strains present in the participants of the MicroAML study (Figure 2D). To further characterize the strains, the supernatant of the 5 *L. eligens* strains were incubated with C2C12 myotubes in the presence of dexamethasone, which established that all strains had anti-atrophic effects (Figure 2E). Considering these results, the subsequent experiments were performed using the supernatant of *L. eligens* CLA-AA-H260, as this strain is representative of the strains found in the participants of the MicroAML study, it does not include putative AMR genes, and grows well and reproducibly.

**Figure 2.**
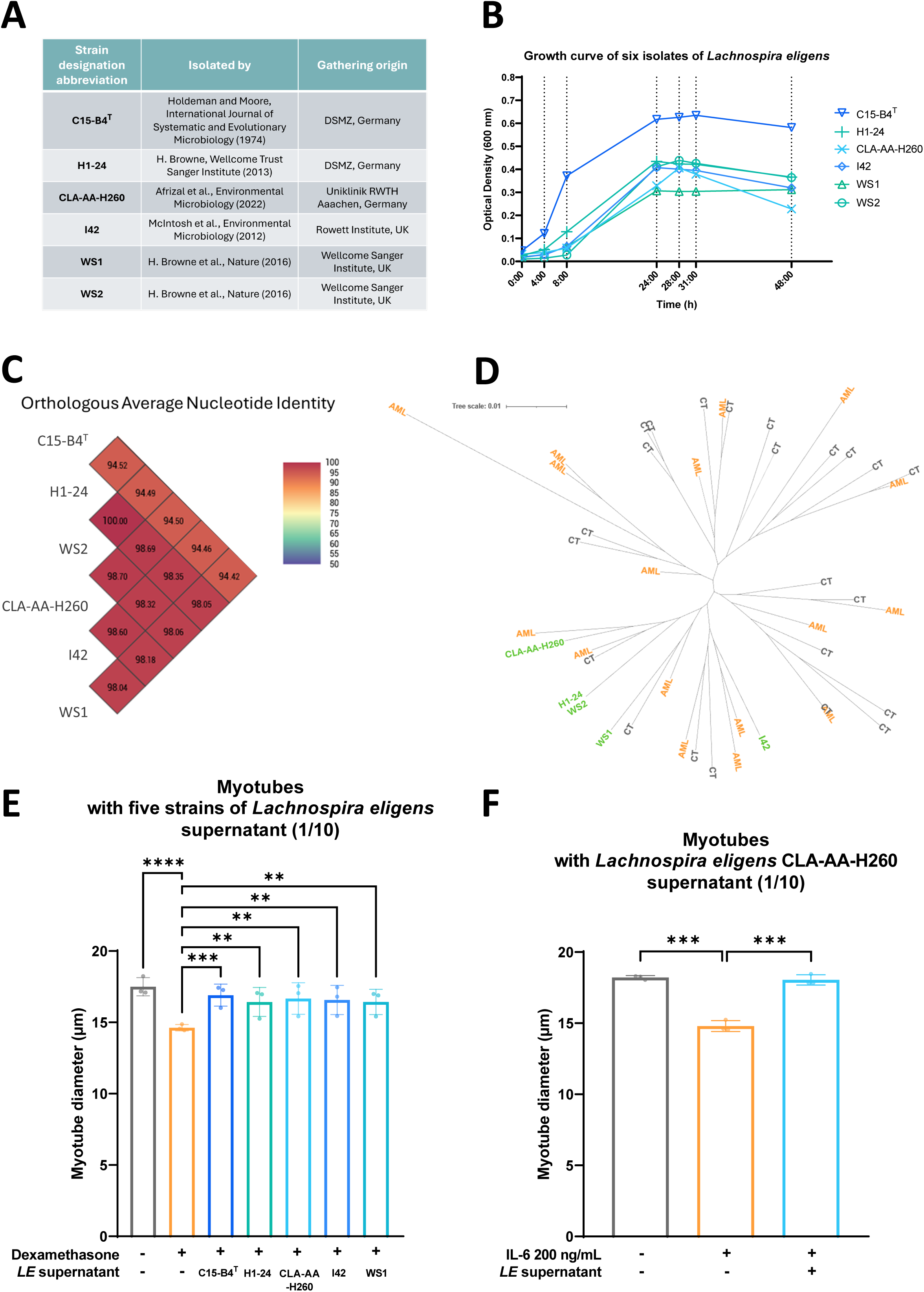
The anti-atrophic effect of *Lachnospira eligens* is conserved between strains and the supernatant of *L. eligens* CLA-AA-H260 counteracts IL-6-induced myotube atrophy. (A) Strain designation abbreviation, isolation and gathering origin of the 6 *L. eligens* isolates. (B) Growth curve over 48 hours of the 6 *L. eligens* isolates. Optical density was measured at 600 nm at the timepoints indicated by a vertical dotted line. Data are presented as mean. (C) Genome comparison of the 6 *L. eligens* isolates by performing the overall genome relatedness index (OGRI) OrthoANI. (D) Phylogenetic tree of the 5 *L. eligens* isolates present in the same Species Genome Bin (SGB 5082) and the *L. eligens* strains present in the microbiota of the participants of the MicroAML study. *L. eligens* strains are shown in green, treatment-naive Acute myeloid leukemia patients in orange and healthy individuals in grey. (E) Comparison of myotube diameter between myotubes incubated with or without 1 µM of dexamethasone and with or without 1/10 diluted supernatant of the 5 *L. eligens* strains. Data are presented as mean ± SD, representative of 3 independent experiments performed in triplicates (N = 3, n = 3). Repeated Measures One-Way ANOVA (p=0.0004) followed by Dunnett’s multiple comparisons test, **p<0.01, ***p<0.001, ****p<0.0001. (F) Comparison of myotube diameter between myotubes incubated with or without 200 ng/mL of interleukin-6 (IL-6) and with or without 1/10 diluted *L. eligens* CLA-AA-H260 supernatant. Data are presented as mean ± SD, representative of 3 independent experiments performed in triplicates (N = 3, n = 3). Repeated Measures One-Way ANOVA (p=0.0002) followed by Dunnett’s multiple comparisons test, ***p<0.001. Abbreviation : *LE* (*Lachnospira eligens*).

### The supernatant of *L. eligens* CLA-AA-H260 counteracts IL-6-induced myotube atrophy

The anti-atrophic effect of *L. eligens* supernatant was evaluated in a different myotube atrophy model, in which C2C12 myotubes were incubated for 48 hours with IL-6 simultaneously or not with *L. eligens* CLA-AA-H260 supernatant. IL-6 induced a significant decrease in myotube diameter which was counteracted by *L. eligens* CLA-AA-H260 supernatant (Figure 2F).

### The anti-atrophic effect of *L. eligens* supernatant is mediated by small non-peptide metabolites

To identify the nature of the active compounds in the culture supernatant of *L. eligens*, an activity-guided biochemical fractionation pipeline was carried out (Figure 3A). First, the supernatant of *L. eligens* CLA-AA-H260 maintained its anti-atrophic effect when filtered at 10, 3 and 1 kDa (Figure 3B). Then, since it has been shown that peptides could have an impact on C2C12 myotubes [62], peptide exclusion experiments were performed and revealed that the supernatant of *L. eligens* CLA-AA-H260 filtered at 1 kDa maintained its anti-atrophic effect when its proteins were denatured by heat or precipitated with ice-cold acetone (Figure 3C). In parallel, an extinction experiment was also performed (Figure 3D). This set of experiments identifies the active compounds as non-peptide metabolites smaller than 1 kDa which remained active after 50 times dilution but not active after 100 times dilution.

**Figure 3.**
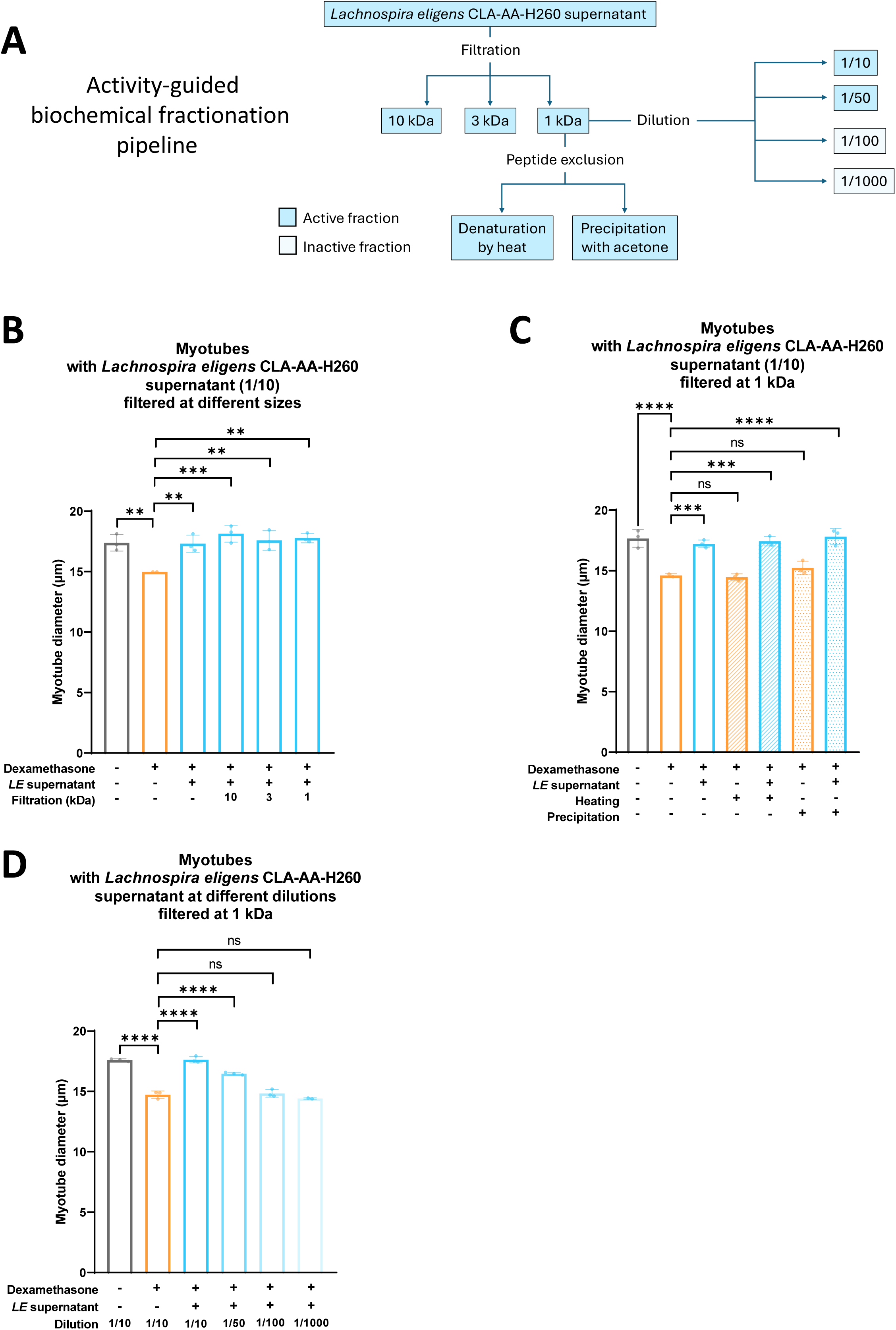
The anti-atrophic effect of *Lachnospira eligens* supernatant is mediated by small non-peptide metabolites. (A) Activity-guided biochemical fractionation pipeline diagram. (B) Filtration experiment. Comparison of myotube diameter between myotubes incubated with or without 1 µM of dexamethasone and with or without 1/10 diluted *L. eligens* CLA-AA-H260 supernatant filtered at 10, 3 and 1 kDa. Data are presented as mean ± SD, representative of 1 independent experiment (N = 1, n = 2-3). One-Way ANOVA (p=0.0043) followed by Dunnett’s multiple comparisons test, **p<0.01, ***p<0.001. (C) Peptide exclusion experiment. Comparison of myotube diameter between myotubes incubated with or without 1 µM of dexamethasone and with or without 1/10 diluted *L. eligens* CLA-AA-H260 supernatant filtered at 1 kDa. Peptides from the NCM and *L. eligens* CLA-AA-H260 supernatant were denatured by heat or precipitated with ice-cold acetone. Data are presented as mean ± SD, representative of 1 independent experiment (N = 1, n = 2-3). One-Way ANOVA (p<0.0001) followed by Dunnett’s multiple comparisons test, **p<0.01, ***p<0.001. (D) Extinction by dilution experiment. Comparison of myotube diameter between myotubes incubated with or without 1 µM of dexamethasone and with or without 1/10, 1/50, 1/100 or 1/1000 diluted *L. eligens* CLA-AA-H260 supernatant filtered at 1 kDa. Data are presented as mean ± SD, representative of 1 independent experiment (N = 1, n = 2-3). One-Way ANOVA (p<0.0001) followed by Dunnett’s multiple comparisons test, ***p<0.001, ****p<0.0001. Abbreviation : *LE* (*Lachnospira eligens*).

Of note, to evaluate if the anti-atrophic effect of the supernatant of *L. eligens* CLA-AA-H260 was effective once muscle atrophy was established, myotubes were incubated 48 hours with dexamethasone to induce an atrophy, followed by an incubation with *L. eligens* CLA-AA-H260 supernatant or NCM (Supplemental Figure 3). *L. eligens* CLA-AA-H260 supernatant did not show a curative effect (Supplemental Figure 3).

### Mixed acid fermentation end-products mediate the anti-atrophic effect of *L. eligens* supernatant

The supernatant of the 5 *L. eligens* strains tested showed an anti-atrophic effect, suggesting one or several common active compound(s). To identify these non-peptidic metabolites smaller than 1 kDa, the supernatant of the 5 *L. eligens* strains were analyzed by ^1^H-NMR metabolomics (Figure 4A). All the strains display a similar pattern of metabolites (Figure 4B), including acetate, formate, lactate and ethanol, which are end-products of mixed acid fermentation (MAF) (Figure 4C).

**Figure 4.**
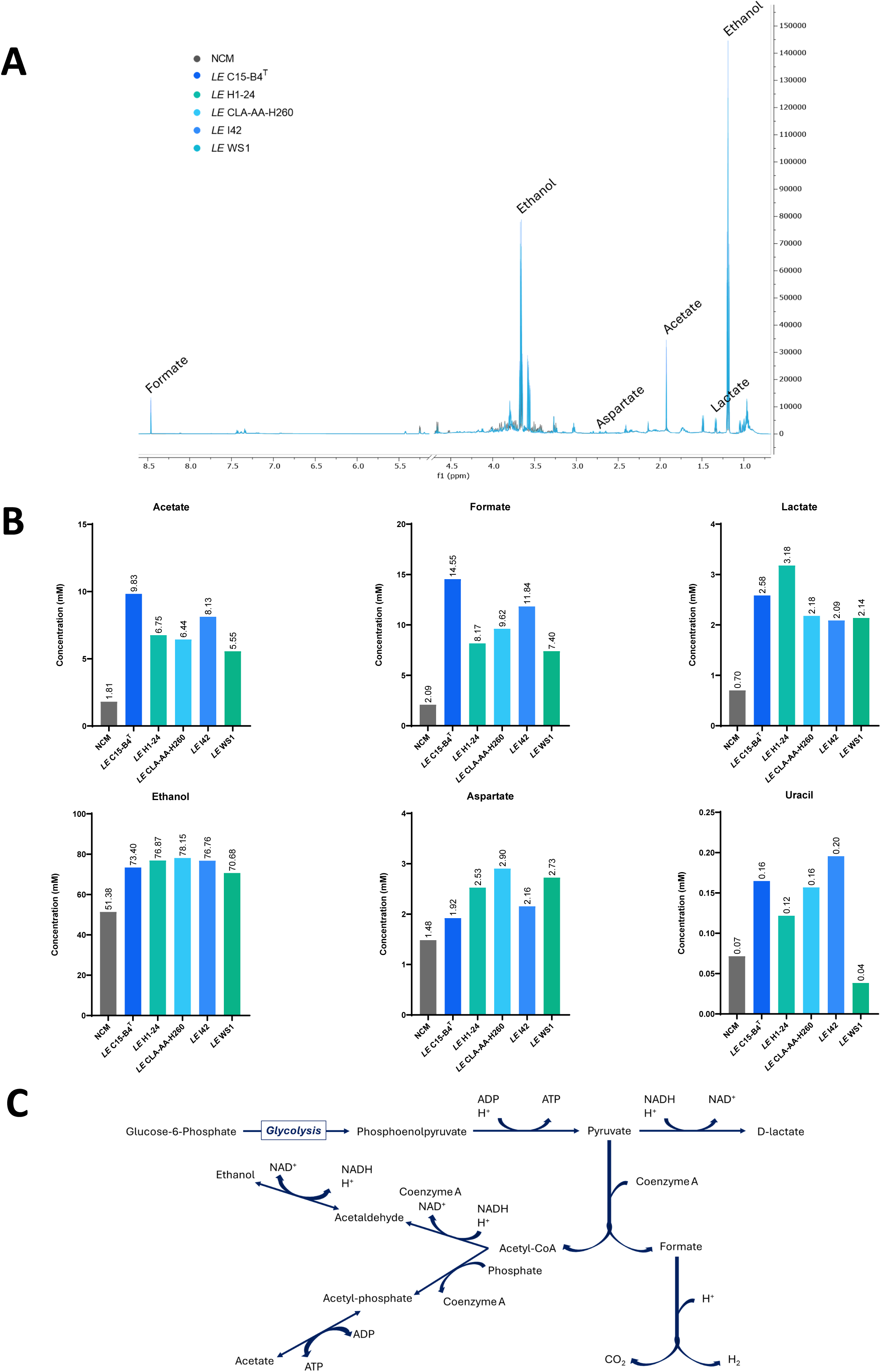
*Lachnospira eligens* strains perform Mixed Acid Fermentation (MAF). (A) Superimposed ^1^H-NMR spectra obtained from non-conditioned medium (NCM) and culture supernatant of the 5 *L. eligens* strains. Uracil could not be visually indicated in the spectra due to its low intensity. (B) Concentration (mM) of metabolites produced by the 5 *L. eligens* strains as well as their concentration in NCM, measured by ^1^H-NMR metabolomics. (C) Diagram of MAF metabolic pathway performed by *L. eligens*. Abbreviation : *LE* (*Lachnospira eligens*).

To determine the importance of MAF, the supernatant of genetically close and distant bacterial species performing MAF were incubated with myotubes in the presence of dexamethasone (Figure 5A). *Lachnospira hominis* CLA-JM-H89B and *Lachnospira hominis* CLA-JM-H10 were chosen as genetically close strains and the commensal strain *Escherichia coli* HS was chosen as a genetically distant strain. Their culture supernatant significantly increased myotube diameter of dexamethasone-treated myotubes (Figure 5A) and contained the end-products of MAF (Figure 5B).

**Figure 5.**
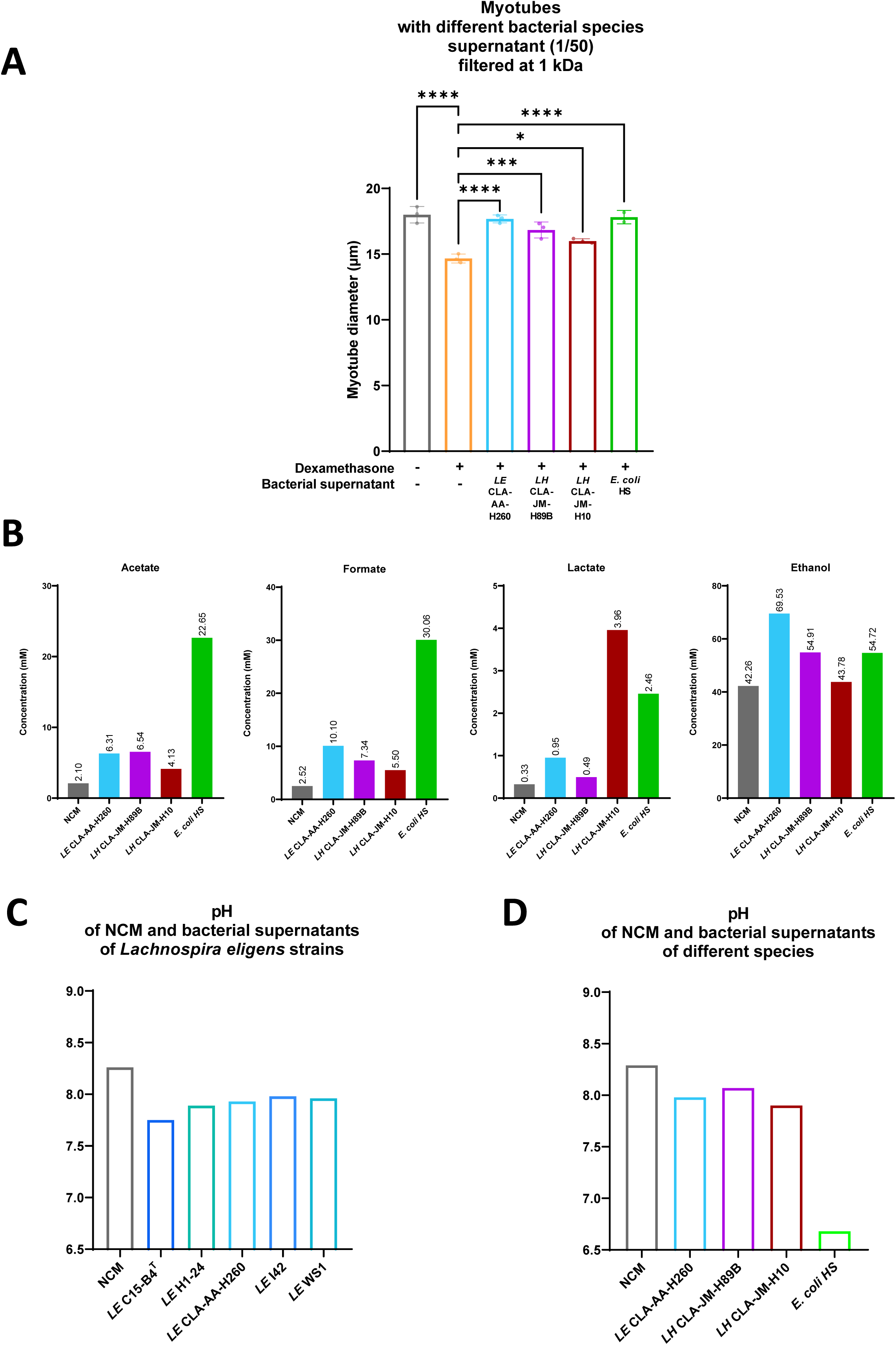
The supernatant of other bacteria performing Mixed Acid Fermentation (MAF) exert an anti-atrophic effect. (A) Comparison of myotube diameter between myotubes incubated with or without 1 µM of dexamethasone and with or without 1/50 diluted and filtered at 1 kDa supernatant of *Lachnospira eligens (LE)* CLA-AA-H260, *Lachnospira hominis (LH)* CLA-JM-H89B, *Lachnospira hominis (LH)* CLA-JM-H10 or *Escherichia coli* HS. Data are presented as mean ± SD, representative of 1 independent experiment (N = 1, n = 2-3). One-Way ANOVA (p<0.0001) followed by Dunnett’s multiple comparisons test, *p<0.05, ***p<0.001, ****p<0.0001. (B) Concentration (mM) of acetate, formate, lactate and ethanol in non-conditioned medium (NCM) and in the culture supernatant of *Lachnospira eligens (LE)* CLA-AA-H260*, Lachnospira hominis (LH)* CLA-JM-H89B*, Lachnospira hominis (LH)* CLA-JM-H10 *and Escherichia coli* HS during MAF, measured by ^1^H-NMR metabolomics. (C) pH of NCM and culture supernatants of the 5 *L. eligens (LE)* strains. (D) pH of NCM and culture supernatant of *Lachnospira eligens (LE)* CLA-AA-H260*, Lachnospira hominis (LH)* CLA-JM-H89B*, Lachnospira hominis (LH)* CLA-JM-H10 *and Escherichia coli* HS.

Interestingly, in accordance with the production of acids during MAF, the pH of the bacterial supernatants was lower than the pH of the NCM (Figure 5C and 5D), with the supernatant of *E. coli* HS exhibiting the lowest pH and the highest concentration of acetate and formate.

Next, to determine whether the acids produced during MAF were the effectors of the anti-atrophic effect observed in dexamethasone-induced atrophied myotubes, the supernatant of *L. eligens* CLA-AA-H260 and *E. coli* HS as well as NCM were filtered at 1 kDa then run through acid-binding columns to extract carboxylic and other weak acids. This approach removed acetate, formate and lactate (Figure 6A and Supplemental Figure 4). As hypothesized, the acid-depleted supernatants of *L. eligens* CLA-AA-H260 and *E. coli* HS did not counteract myotube atrophy (Figure 6B). In addition, when myotubes were incubated with acetate, formate or D-lactate diluted in NCM at the concentration found upon MAF by *L. eligens*, the acids counteracted the atrophy induced by dexamethasone (Figure 6C).

**Figure 6.**
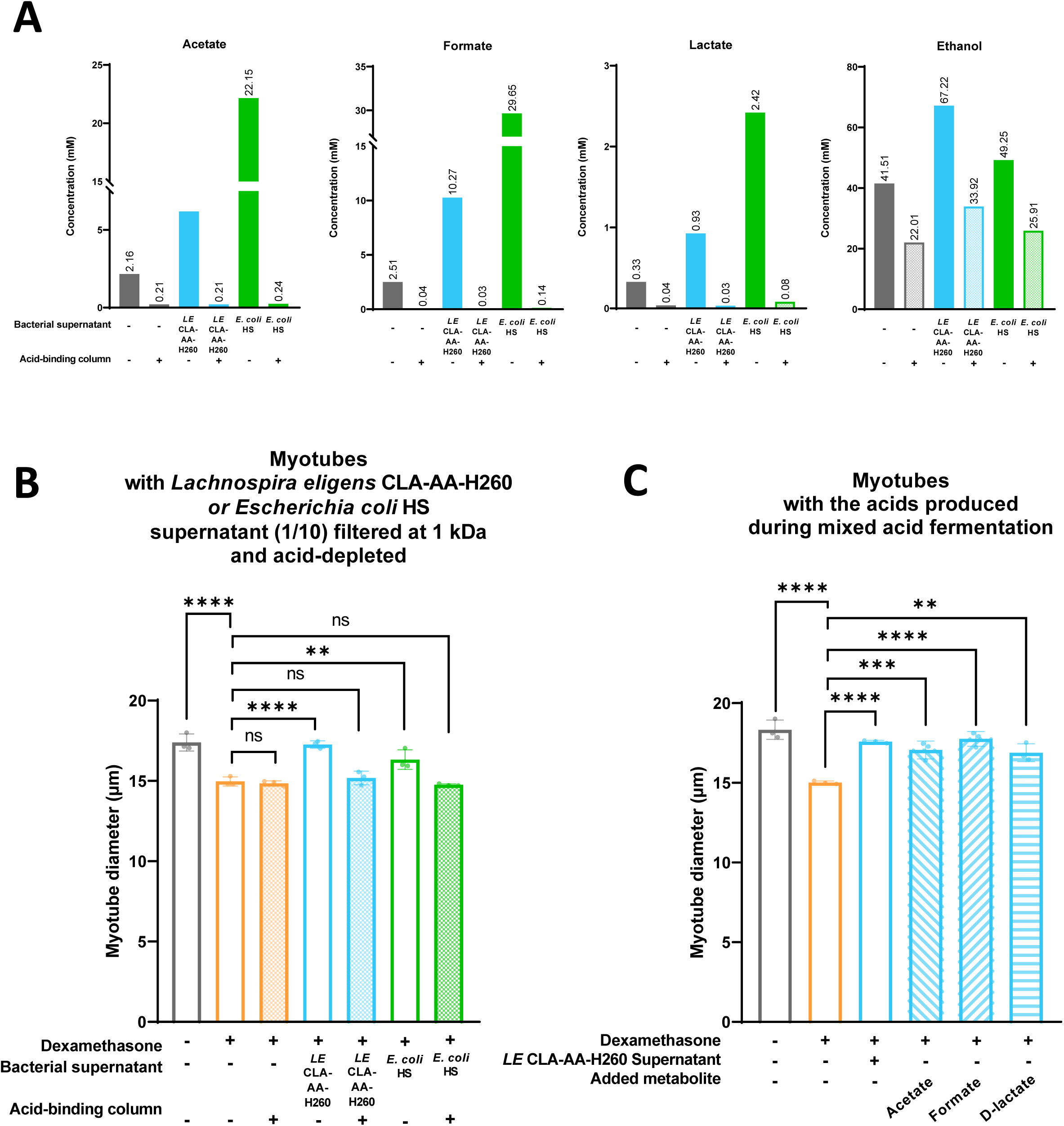
Mixed acid fermentation end-products mediate the anti-atrophic effect of *Lachnospira eligens* supernatant. (A) Concentration (mM) of acetate, formate, lactate and ethanol in non-conditioned medium (NCM) and in the culture supernatant of *Lachnospira eligens (LE)* CLA-AA-H260 *and Escherichia coli* HS after they had been run or not through acid-binding columns. Concentrations were measured by ^1^H-NMR metabolomics. (B) Comparison of myotube diameter between myotubes incubated with or without 1 µM of dexamethasone and with or without 1/10 diluted and filtered at 1 kDa supernatant of *Lachnospira eligens (LE)* CLA-AA-H260 *or Escherichia coli* HS after they had been run or not through acid-binding columns. Data are presented as mean ± SD, representative of 1 independent experiment performed in triplicates (N = 1, n = 3). One-Way ANOVA (p<0.0001) followed by Dunnett’s multiple comparisons test, **p<0.01, ****p<0.0001. (C) Comparison of myotube diameter between myotubes incubated with or without 1 µM of dexamethasone and with or without 1/10 diluted supernatant of *Lachnospira eligens (LE)* CLA-AA-H260 or NCM with acetate, formate or D-lactate at 7.47 mM, 10.51 mM and 2.55 mM respectively. Data are presented as mean ± SD, representative of 1 independent experiment performed in triplicates (N = 1, n = 3). One-Way ANOVA (p<0.0001) followed by Dunnett’s multiple comparisons test, **p<0.01, ***p<0.001, ****p<0.0001.

### The anti-atrophic effect of *L. eligens* occurs independently of a modulation of the transcriptome and of the net consumption of MAF end-products

The next step aimed to unravel the mechanism of action on the host side. To do so, we analyzed by RNA sequencing the transcriptome of C2C12 myotubes incubated or not with the supernatant of *L. eligens* CLA-AA-H260 filtered at 1 kDa, in the presence or not of dexamethasone, for 48 hours. PCA revealed a clear separation between samples treated or not with dexamethasone whereas no separation was observed between samples treated or not with the supernatant of *L. eligens* CLA-AA-H260 in the presence of dexamethasone (Figure 7A). The latter was confirmed by the presence of only one differentially expressed gene between groups treated or not with the supernatant of *L. eligens* CLA-AA-H260 in the presence of dexamethasone (Figure 7B), namely *Serpina3n* (p-adjusted < 0.05). *Serpina3n* expression, a biomarker of muscle atrophy [63, 64], was increased in dexamethasone-induced atrophied myotubes compared to control, and this increase was partially counteracted by the supernatant of *L. eligens* CLA-AA-H260 (Figure 7C). These results were validated by qPCR (Supplementary Figures 5A and 5B).

**Figure 7.**
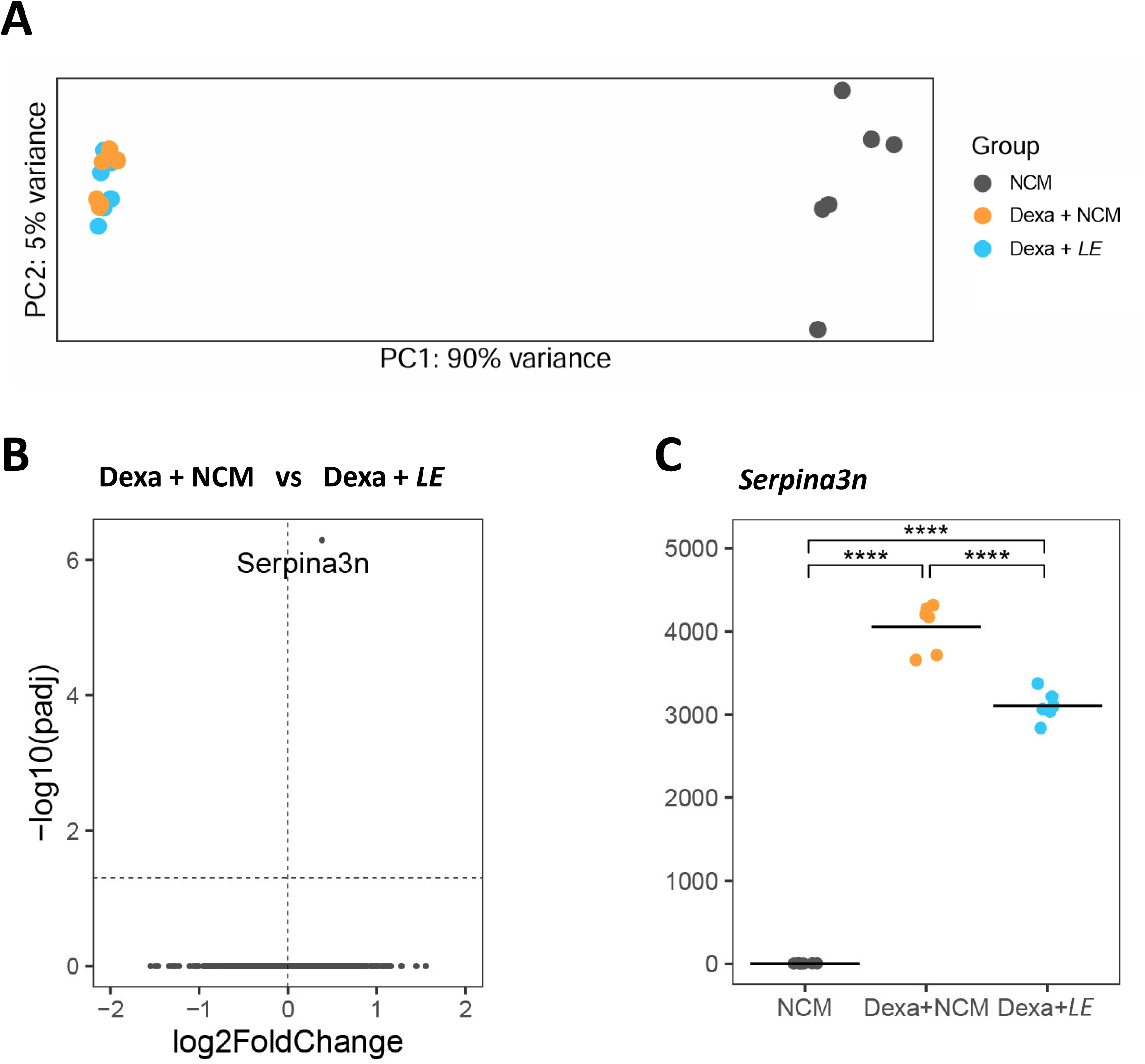
The anti-atrophic effect of *Lachnospira eligens* CLA-AA-H260 occurs independently of a modulation of the transcriptome. (A) Principal Component Analysis (PCA) performed on RNAseq variance-stabilized counts. Pool of 2 independent experiments performed in triplicates. (B) Volcano plot showing the result of a differentially expression analysis between myotubes incubated with dexamethasone and 1/10 diluted and 1 kDa filtered *L. eligens* CLA-AA-H260 supernatant (Dexa + *LE*) and myotubes incubated with dexamethasone and non-conditioned medium (Dexa + NCM). Genes with an adjusted p-value < 0.05 and a log2 fold change > 1 were considered differentially expressed. (C) RNAseq normalized counts of *Serpina3n* (DESeq2 normalization method). ****p<0.0001.

In parallel, the metabolic net flux of MAF acid end-products and glucose was evaluated by analyzing their concentration in wells with or without myotubes after 48 hours by ^1^H-NMR metabolomics. Lactate and glucose were clearly secreted and consumed respectively, with a lower flux upon dexamethasone, with or without the supernatant of *L. eligens*. No net consumption of acetate and formate was observed, independently of the condition (Supplemental Figure 6).

### Shifting from anoxic to oxic culture conditions prevents the anti-atrophic effect of *E. coli* HS supernatant

To validate that the anoxic culture conditions and the MAF performed by some bacteria in this oxygen-depleted environment are necessary to exert the anti-atrophic effect observed in atrophied C2C12 myotubes, we decided to culture a bacterium in anoxic and oxic conditions. *L .eligens* is an obligate anaerobic bacterium requiring anoxic conditions to grow. In contrast, *E. coli* HS is a facultative anaerobe that can grow both in anoxic and oxic conditions. The supernatants of *E. coli* HS cultured in anoxic and oxic conditions were therefore used for subsequent experiments. At this stage, a defined culture medium containing glucose, cystine, glycerol and salts was used to fully control its composition, in the perspective of an *in vivo* administration and a potential industrial production for clinical purposes. As hypothesized, the supernatant of *E. coli* HS cultured in anoxic conditions counteracted the atrophying effect of dexamethasone whereas the supernatant of *E. coli* HS cultured in oxic conditions did not (Figure 8A). The supernatant of *E. coli* HS cultured in anoxic conditions contained the acid end-products of MAF, whereas the supernatant of *E. coli* HS cultured in oxic conditions did not (Figure 8C), which is in accordance with the former being more acidic than the latter (Figure 8B).

**Figure 8.**
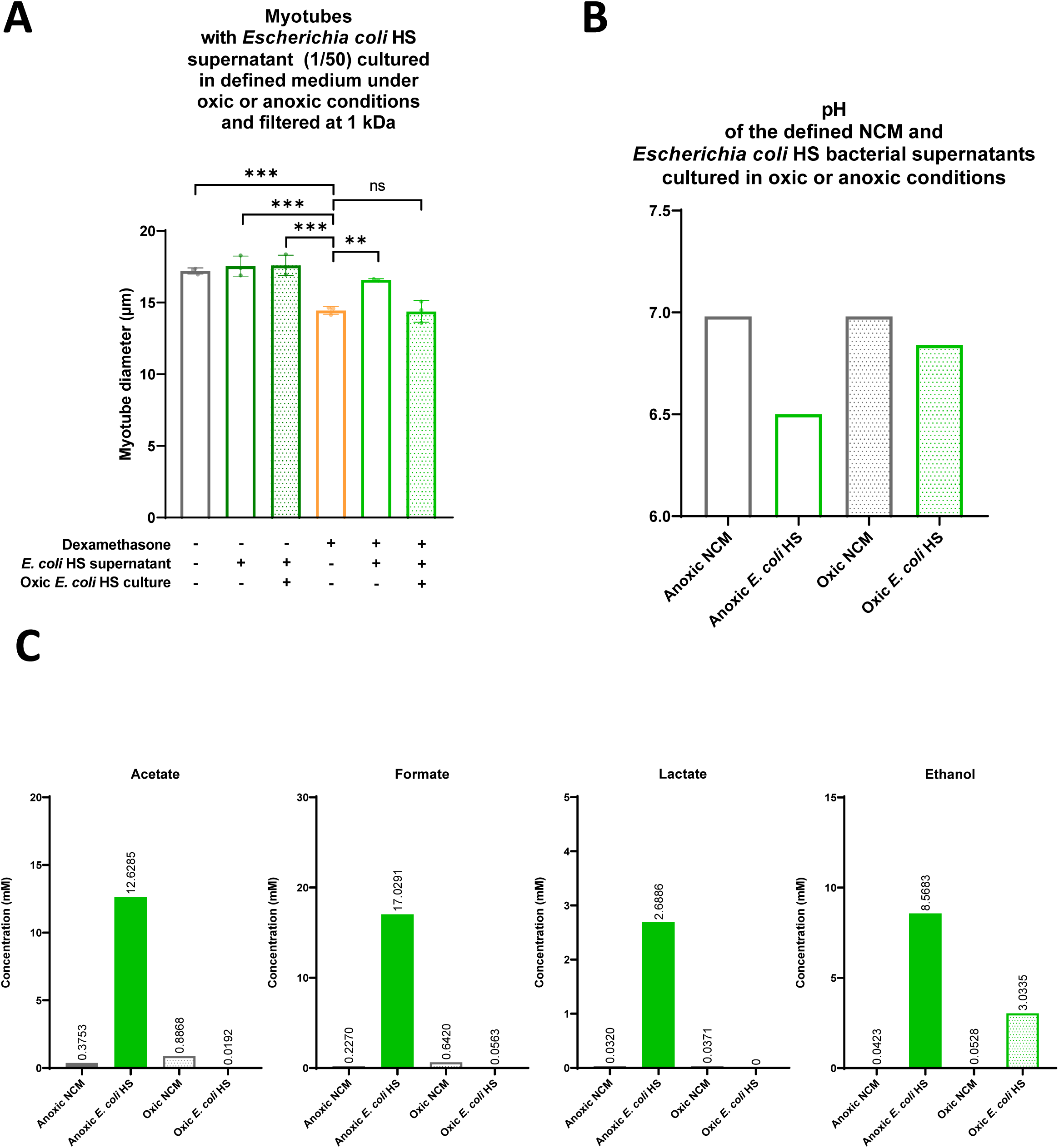
Anoxic culture conditions are necessary for the anti-atrophic effect of *Escherichia coli* HS supernatant. (A) Comparison of myotube diameter between myotubes incubated with or without 1 µM of dexamethasone and with or without 1/50 diluted and 1 kDa filtered supernatant of *E. coli* HS cultured in defined medium in oxic or anoxic conditions. Data are presented as mean ± SD, representative of 1 independent experiment (N = 1, n = 2-3). One-Way ANOVA (p<0.0001) followed by Dunnett’s multiple comparisons test, **p<0.01, ***p<0.001. (B) pH of the defined non-conditioned medium (NCM) and culture supernatants of *E. coli* HS cultured in defined medium under oxic or anoxic conditions. (C) Concentration (mM) of acetate, formate, lactate and ethanol in NCM and in the culture supernatants of *E. coli* HS cultured in defined medium under oxic or anoxic conditions. Concentrations were measured by ^1^H-NMR metabolomics. Abbreviation : NCM (non-conditioned medium).

### Supplementation with *E. coli* HS anoxic supernatant does not prevent muscle atrophy and weakness in leukemic mice

With the aim of finding a treatment to tackle muscle atrophy in cancer cachexia, the effects of the supernatants of *E. coli* HS cultured in anoxic and oxic conditions were tested in the BaF model (Figure 9A). The supernatants of *E. coli* HS were selected for this *in vivo* experiment, rather than the supernatant of *L. eligens*, to allow for the inclusion of two negative controls, namely the non-conditioned medium and the supernatant of *E. coli* HS in oxic conditions The solutions were concentrated to increase the concentration of MAF acid end-products and then filtered at 1 kDa. Their MAF end-products levels, or lack thereof, were validated using ^1^H-NMR metabolomics (Supplemental Figure 7).

**Figure 9.**
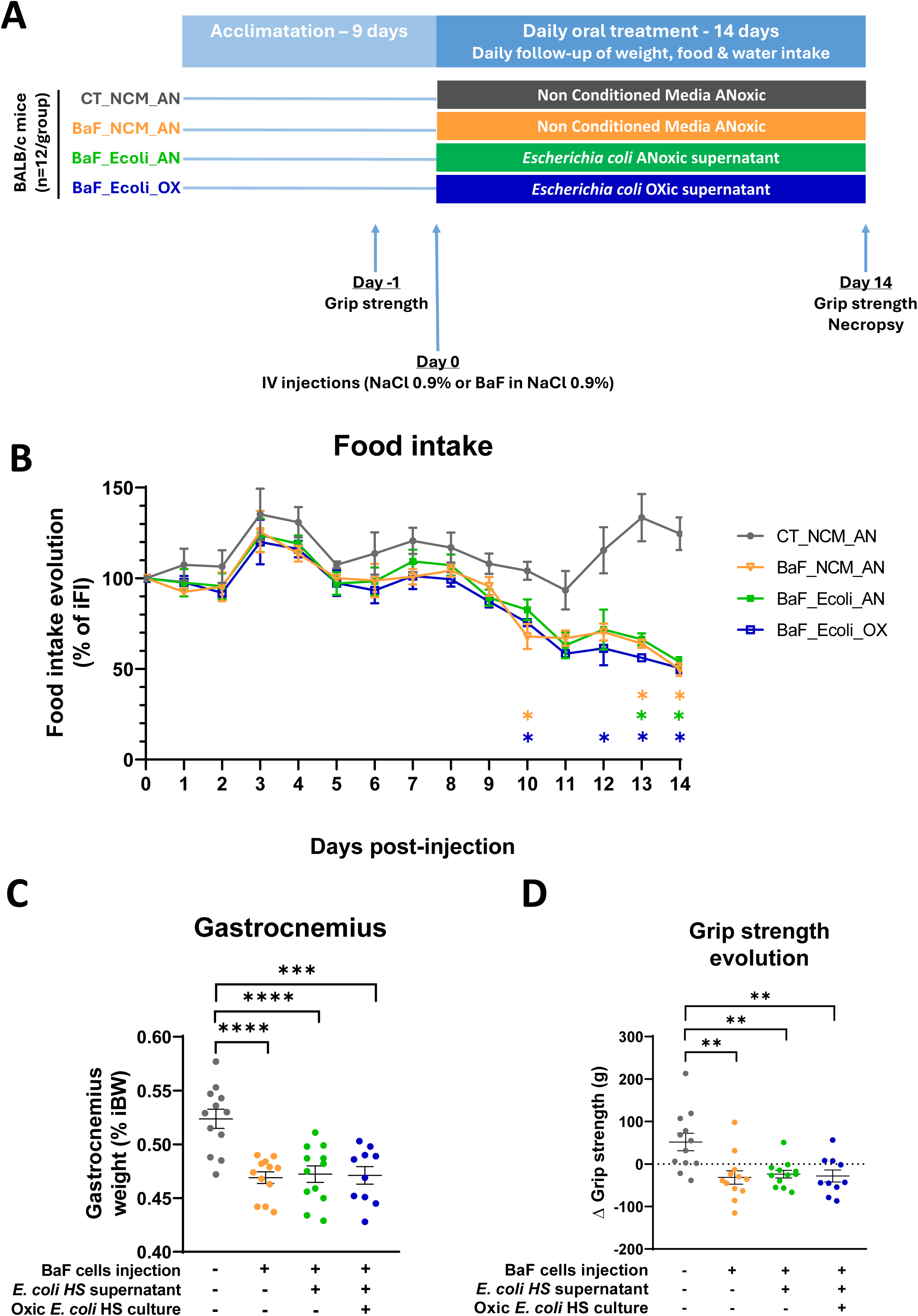
Supplementation with *Escherichia coli* HS anoxic supernatant does not prevent muscle atrophy and weakness in leukemic mice. (A) Diagram of the experimental design for the *in vivo* experiment. (B) Evolution of food intake in percentage of initial food intake. Data are presented as mean ± SEM. Mixed-effects model with the Geisser-Greenhouse correction was performed followed by Tukey’s multiple comparisons test, p=ns are not shown, significant comparisons (*p<0.05) were pairwise comparisons of the group CT_NMC_AN vs BaF_NCM_AN, vs BaF_Ecoli_AN and vs BaF_Ecoli_OX. (C) Gastrocnemius muscle weight in percentage of initial body weight. Data are presented as mean ± SEM. One-Way ANOVA (p<0.0001) was performed followed by Tukey’s multiple comparisons test, ***p<0.001, ****p<0.0001. (D) Grip strength evolution between the start and the end of the experiment. Data are presented as mean ± SEM. One-Way ANOVA (p<0.0008) was performed followed by Tukey’s multiple comparisons test, **p<0.01.

Cachexia was observed in the 3 groups injected with leukemic cells (BaF_NCM_AN, BaF_Ecoli_AN and BaF_Ecoli_OX) compared to control (CT_NCM_AN), as shown by a significant decrease in food intake on the last two days (Figure 9B) as well as a significant decrease in gastrocnemius weight on the day of necropsy (Figure 9C) and a significant decrease in grip strength between the start and the end of the experiment (Figure 9D). No significant effect of the administration of the supernatant of *E. coli* HS cultured in anoxic conditions was observed on those parameters. As reported before [27], despite the presence of cachexia, body weight did not significantly decrease in the 3 leukemic groups compared to control (Supplemental Figure 8A). This can be explained by a significant increase of the spleen and liver weights due to leukemic cells infiltration (Supplemental Figure 8B and 8C). No significant effect of the administration of the supernatant of *E. coli* HS cultured in anoxic conditions was observed on those parameters. In accordance with the phenotypic observations, skeletal muscle atrogenes and *Serpina3n* were upregulated to the same extent in the gastrocnemius and soleus of the 3 leukemic groups (Supplemental Figure 9A and 9B). In the jejunum, the 3 leukemic groups showed a similar decrease in tight-junction integrity (*Ocln* and *Tjp1*) and mucosal inflammation (*Il1b*) but preserved protection (*Muc2*) (Supplemental Figure 9C), whereas in the ileum, they showed a comparable decrease in mucosal inflammation (*Il1b*) and protection (*Muc2*) but a preserved tight-junction integrity (*Ocln* and *Tjp1*) (Supplemental Figure 9D). To understand why the administration of the supernatant of *E. coli* HS cultured in anoxic conditions, containing MAF acid end-products, did not counteract muscle atrophy and weakness, the systemic blood levels of acetate, formate and lactate on the day of necropsy were measured by ^1^H-NMR metabolomics. Acetate and formate levels did not vary between groups whereas lactate levels increased in the BaF_NCM_AN and BaF_Ecoli_AN groups and showed a tendency (p=0.0520) to increase in the BaF_Ecoli_OX group compared to the CT_NCM_AN healthy mice group (Supplemental Figure 10).

### *L. eligens* levels are associated with low muscle strength, low blood acetate levels and high relative abundances of fecal oral bacteria and facultative anaerobes in patients

In the MicroAML cohort, where the relative abundance of *L. eligens* was reduced in treatment-naive patients with AML, muscle strength was reduced and associated with reduced blood levels of acetate (Figure 10A) while the other MAF end-products formate and lactate were not reduced [4]. Re-analysis of the metagenomics data of the feces of these patients confirmed an increase in oral bacteria in the feces of patients with AML, suggesting an increase in O_2_ levels in the gut. Considering this result and given the importance of anoxic conditions to foster MAF, the community oxygen preference profile was explored. An increase in the relative abundance of bacteria capable of performing aerobic respiration and more specifically facultative anaerobes was observed in treatment-naive patients with AML. Finally, the link between *L. eligens* and muscle strength was explored in MASLD, since its association with sarcopenia has been reported, [9, 10]. Patients with MASLD firstly experience sarcopenia, which is often associated with obesity [9], while cachexia may occur at the end-stage of disease, in the presence of cirrhosis [65]. Interestingly, the exploration of the INSYTE cohort [37] showed that grip strength tended to increase in patients with *L. eligens* levels higher than median (n=44) compared to patients with *L. eligens* levels lower than median (n=45) (p=0.1062) (Figure 10D).

**Figure 10.**
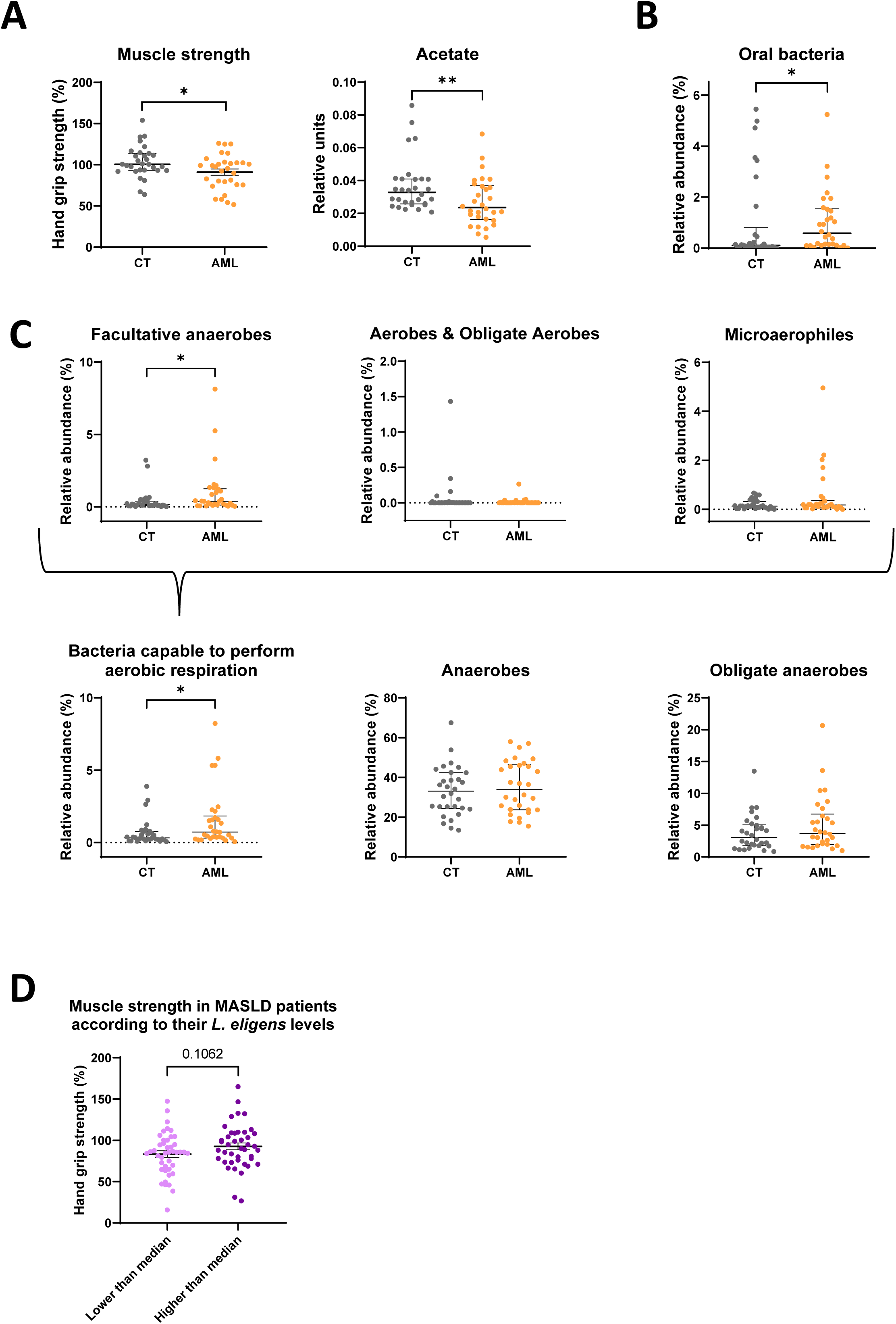
*Lachnospira eligens* levels are associated with low muscle strength, low blood acetate levels and high relative abundances of fecal oral bacteria and facultative anaerobes in patients. (A) Muscle strength and blood acetate levels in treatment-naive patients with acute myeloid leukemia (AML) compared to healthy individuals. Results have been previously published in Pötgens et *al.*, *Haematologica* 2024. (B) Oral species abundance in the feces of patients with AML compared to healthy individuals. (C) Community oxygen preference profile in patients with AML compared to healthy individuals. (D) Muscle strength of Metabolic Dysfunction-Associated Steatotic Liver Disease (MASLD) patients with levels of *L. eligens* higher than median (n=44) compared to patients with *L. eligens* levels lower than median (n=45). Normally-distributed data are presented as mean ± SEM. Student t test was performed (*p<0.05). Non-normally distributed data are presented as median ± IQR. Mann-Whitney U test was performed ((*p<0.05, **p<0.01).

## Discussion

Our work contributes to establishing the clinical relevance of studying *Lachnospira eligens*. Indeed, *L. eligens* abundance was found to be reduced in 2 independent cohorts of treatment-naive patients with AML. In the MicroAML cohort, *L. eligens* abundance was strongly correlated with muscle strength, an important hallmark of cancer cachexia [4]. In agreement with this observation, *L. eligens* was reduced in cachectic *versus* non-cachectic patients with metastatic cancer [66]. *L. eligens* abundance has also been shown to be reduced in postmenopausal breast cancer patients [67] and in esophageal squamous cell carcinoma patients [68]. Along these lines, higher abundances of *L. eligens* were associated with a decreased colorectal cancer (CRC) probability [69], consistent with the active patent for the use of *L. eligens* strains in the prevention and treatment of CRC-related diseases [70].

Beyond the context of cancer, *L. eligens* abundance was also positively associated with grip strength in a cohort of older adults [71]. Reduced *L. eligens* abundance was also reported in patients with Cushing’s syndrome [72], an endocrine disease caused by prolonged exposure to high levels of glucocorticosteroids medications or cortisol, leading to reduced muscle strength [73]. Consistently, we also report here that patients with MASLD with a low relative abundance of *L. eligens* tend to display lower muscle strength than patients with high relative abundances of *L. eligens*. Taken together, these results corroborate that *L. eligens* may be relevant for muscle health in cancer, sarcopenia, MASLD and other disorders affecting muscle health.

In the gut, mixed acid fermentation (MAF) is performed by bacteria which have different oxygen tolerance such as obligate anaerobes like *L. eligens* or facultative anaerobes like *E. coli*. In the presence of O_2_, facultative anaerobes can perform aerobic respiration, which has a higher ATP yield than anaerobic respiration and fermentation. In the presence of O_2_ in the intestine, the growth of bacteria that have the metabolic flexibility to perform aerobic respiration is therefore enhanced over the others and confers them a competitive advantage [74]. Likewise, in the presence of other electron acceptors such as sulfate (SO_4_^2-^), nitrate (NO_3_^-^), ferric iron (Fe_3_^+^) and fumarate, the growth of bacteria performing anaerobic respiration is enhanced. In those situations, bacteria which rely solely on fermentation are outpaced and their relative abundance decreases.

The type strain of *L. eligens* was identified as a low-tolerance taxa to H₂O₂ and O₂ stress [75]. More broadly, oxidative stress induced by H₂O₂ and O₂ consistently altered gut microbial communities, depleting strict anaerobes, particularly butyrate producers and enriching facultative or aerotolerant taxa [75]. Interestingly, it has been proposed that consumption of the available O₂ by facultative anaerobes makes the intestinal environment more anaerobic and thus promote the stability of the predominantly anaerobic microbes [76], contributing to homeostasis. Another explanation is also emerging, namely that O_2_ and nitrate levels in the intestine (and therefore electron acceptor availability) are primarily driven by the host through the regulation of intestinal epithelial hypoxia and inducible nitric oxide synthase (iNOS) synthesis, rather than by microbial consumption [76]. In line with this framework, in diseases caused by enteric pathogens, a bloom of facultative anaerobes has been observed in the colon and one of the ecological drivers of this dysbiosis would be an increased availability of host-derived electron acceptors favoring respiration [77]. Such phenomenon could be extrapolated to noninfectious human conditions characterized by the same dysbiotic microbial signature [77] and has been previously reported in a mouse model of cancer cachexia by our team [78] as well as in mouse models of colitis [79, 80], colitis-associated CRC [81] and graft-*versus*-host disease [82]. In line with the increase of oxygen levels in the intestine and of bacteria performing aerobic respiration in the gut, increased levels of microbes of oral origin have been reported in the gut microbiota of patients with several diseases [83, 84].

This framework is consistent with our results showing that (1) bacteria performing aerobic respiration, and more specifically facultative anaerobes, are increased in treatment-naive patients with AML; (2) their gut microbiota displays a pro-oxidative status as shown by alterations in genes catalyzing redox reactions and in fecal metabolome [4]; (3) the relative abundance of oral species increased more than two-fold in patients with AML.

The work of Pitt *et al*. is a convincing example of the links between bloom of pathogens, lack of gut anaerobiosis and low MAF. They showed that the administration of respiring bacterial membrane vesicles (MV) to a mouse model of gut inflammation restored gut anaerobiosis, with reduced *Enterobacteriaceae* and increased anaerobic *Lactobacillaceae*, and ameliorated symptoms such as weight loss and shortening of the colon [85]. Moreover, metatranscriptomic analysis of the microbiome showed a general shift from respiration to fermentation in MV-treated mice, with MAF highlighted as one of the most discriminant pathways [85].

Taken together, these results corroborate the disturbance of electron acceptors levels in the gut of patients with AML and raise the hypothesis that *L. eligens* might be a possible biomarker of disturbed gut O_2_ content, explaining its decreased abundance in several conditions.

During MAF, *L. eligens* produces the acids acetate, formate and D-lactate. Once those acids are absorbed by enterocytes, they are transported to the liver and then released into systemic circulation, where they can reach the muscle. As a cocktail, SCFA administration (60:25:15 for acetate:propionate:butyrate) can improve muscle health, with reports indicating decreased myotube atrophy and atrophy markers induced by dexamethasone and lipopolysaccharides [86, 87] and improved muscle mass and function in a mouse model of sarcopenia [87].

To our knowledge, the effects of each acid individually (acetate, formate or D-lactate) on atrophied myotubes had not been previously documented. In addition, no studies evaluating the effects of formate or D-lactate administration in *in vivo* models of muscle atrophy were found. Acetate has been shown to exerts protective effects on murine skeletal muscle *in vivo*, as its oral administration was shown to reduce atrogene expression and enhance oxidative muscle features in aged rats [88], while also rescuing antibiotic-induced losses in muscle strength and fiber size in young adult mice [89]. Acetate can also be present in the culture supernatant of heterofermentative lactic acid bacteria, for which many studies have reported anti-atrophic effects in dexamethasone atrophied C2C12 myotubes [90–92]. Finally, oral administration of homofermentative lactic acid bacteria in a dexamethasone-induced mouse model of muscle atrophy rescued muscle mass and strength and reduced atrophy markers while increasing fecal acetate levels [93, 94].

Taken together, our findings, as well as the current body of research, support the hypothesis that acetate plays a key role in the benefits conferred by bacterial supernatants preventing muscle atrophy.

To unravel the mechanism of action from the host side, transcriptomics and metabolic net flux of MAF end-products and glucose were performed after treatment with *L. eligens* CLA-AA-H260 supernatant.

Transcriptome analysis pinpointed only one differentially expressed gene, *Serpina3n*. The increased mRNA and protein levels of Serpina3n and its human orthologue SERPINA3 has been convincingly proposed as a biomarker of muscle atrophy [63, 64]. However, no clear functional role has been identified for Serpina3n or its human orthologue in muscle atrophy, outside the context of muscular dystrophy in which Serpina3n has a demonstrated protective function through protease inhibition and stabilization of the sarcolemma [95]. The mechanisms responsible for the induction of Serpina3n or its human orthologue during glucocorticoid-induced atrophy have been previously explored and may involve direct regulation by the glucocorticoid receptor or indirect regulation through FOXO and mTOR [63, 96, 97]. Based on the results of Gueugneau *et al*. [63], the increased expression of *Serpina3n* in muscle treated with glucocorticoids could be considered as a compensatory or protective mechanism to inhibit proteases when protein synthesis is impaired or reduced. Therefore, the rescue of *Serpina3n* expression in dexamethasone-atrophied myotubes incubated with bacterial supernatant containing MAF end-products seems to be the consequence of (or an effect concomitant to) the prevention of myotube atrophy, more than the causal driver.

In parallel, the metabolic net flux of acetate, formate, lactate and glucose was evaluated after 48h of treatment. The consumption of glucose and the secretion of lactate are coherent with the glycolytic metabolism of C2C12 cells. In the presence of dexamethasone, C2C12 cells consumed less glucose, which is in line with previous studies in C2C12 myotubes [98] and in skeletal muscles of rats [99, 100]. While myotube uptake of acetate has been previously reported in primary human and rat myotubes [101, 102], in our experiment, net acetate flux was not reduced. Considering that levels of glucose were still high after 48h of treatment (> 20mM), it can be hypothesized that high glucose availability may have led to a preference for glucose metabolism and therefore prevented the cells from consuming acetate as a source of energy. In addition, a net secretion of formate was observed independently of dexamethasone treatment, which was prevented by *L. eligens* CLA-AA-H260 supernatant, suggesting a decrease in mitochondrial one-carbon metabolism induced by the supernatant. Metabolic tracing of MAF acids would be informative to better define their fate.

Taken together, these results indicate that *L. eligens* CLA-AA-H260 supernatant exerts its effect at the phenotypic level without transcriptome reprogramming and net uptake of MAF acids and led us to the hypothesis that *L. eligens* CLA-AA-H260 supernatant may counteract myotube atrophy through extracellular mechanism(s). Putative mechanisms of action could be extracellular receptor activation such as G protein-coupled receptors (GPCRs). GPR41 (FFAR3) and GPR43 (FFAR2) are well known receptors that are activated by SCFAs, such as acetate (and formate to a much weaker extent) [103, 104]. Activation of GPR41 and GPR43 in rodent myotubes led in both cases to an increased calcium flux [105, 106], indicating the existence of such pathway in myotubes, but the biological role of these pathways in atrophic conditions remains unclear. Based on these data, we postulated that acetate may counteract dexamethasone-induced atrophy through the activation of GPR43 and/or GPR41. This hypothesis could however not be tested in a straightforward manner in our experimental setting as GPR43 antagonists are not active on the mouse isoform and the main antagonist of GPR41, β-hydroxybutyrate, shows weak potency and specificity [107].

The results of the *in vivo* experiment bring a strong limitation to the therapeutic potential of the approach brought forward by this work and raises questions about how to successfully induce a long-lasting increase in the levels of MAF end-products in the blood of mammals. The fact that the oral administration of the bacterial supernatant rich in MAF acid end-products did not counteract muscle atrophy and weakness in leukemic mice may be related to the severity of the phenotype as well as to the lack of increase in the blood levels of MAF end-products. Potential explanations for this lack of increase include a low bioavailability of the acids, a clearance too rapid for them to exert an effect on muscle and insufficient levels of MAF end-products in the supernatant. Information about dosage of individual MAF acids needed to obtain *in vivo* effects on muscle physiology is scarce. In the current work, leukemic mice received an average dose of acetate of 22 mg/kg/day for 14 days, which is below the dosage administered in aged rats (namely 52 mg/kg/day for 19 weeks, 5 days/week) for which a beneficial impact on muscle biomarkers such as atrogenes were reported [88]. The hypothesis of rapid clearance is supported by the observation that a single dose of acetate is eliminated within 120 min in healthy volunteers [108]. To overcome this clearance issue, SCFA are usually administered to rodents through the drinking water. However, such mode of administration was not compatible with the disease model characteristics that include anorexia and reduced water intake. A rapid clearance could be remediated using an osmotic pump enabling subcutaneous continuous infusion [109]. Another alternative that may be considered is the probiotic approach. However, as *L. eligens* does not include any mouse commensal strains (https://microbeatlas.org/landing) and as the gut environment of cachectic mice may disfavor MAF by *L. eligens*, this approach was not selected as a first-line approach. Additional limitations to this work should be acknowledged. Firstly, acid-binding columns removed MAF acid end-products from the supernatant but also decreased by half ethanol and we cannot exclude the elimination of other weak acids. Secondly, ^1^H-NMR metabolomics did not allow to differentiate D-lactate from L-lactate identification. However, this limitation could be compensated by considering knowledge from literature that MAF produces D-lactate. Thirdly, absolute quantification of acetate, formate and lactate in mouse blood could not be achieved due to the interaction of blood albumin with the quantification reference. Fourthly, whole-genome sequencing of the 6 *L. eligens* isolates gathered for this work indicate that the type strain of *L. eligens*, namely *L. eligens* C15-B4^T^, is genetically separated from the other *L. eligens* strains, suggesting that taxonomical classification of these strains may need to be refined. Fifthly, potential indirect beneficial effects of *L. eligens* on muscle health through the promotion of anti-inflammatory cytokines [110] or cross-feeding mechanisms [110–115] were not evaluated in this study.

In conclusion, the reported results provide evidence for an anti-atrophic effect of *L. eligens* supernatant in myotubes, mediated by anoxic MAF end-products, which are small molecules that can enter systemic circulation and could potentially reach the muscle. Further studies would be needed to prove this effect *in vivo*. In addition, our findings corroborate the disturbance of electron acceptors levels in the gut of treatment-naive patients with AML and raises the hypothesis that the oxygen-sensitive obligate anaerobic bacterium *L. eligens* might be a possible biomarker of disturbed gut O_2_ content, explaining its decreased abundance in several conditions. Overall, this work emphasizes the importance of future studies aimed at investigating and therapeutically modulating electron acceptor availability, such as O_2_, in the intestinal environment.

## Data availability statement

Raw RNAseq data can be found on GEO (GEO ID: GSE328252). Bacterial sequencing data can be found in the SRA database : whole-genome sequences of *L. eligens* strains (SRA project ID: PRJNA1457417). Metagenomics and 16S rRNA gene sequences of the MicroAML cohort (SRA project ID: PRJNA813705), 16S rRNA gene sequences of the Wang cohort (SRA project ID: PRJNA746137 for R1 reads, R2 reads were sent upon request) and 16S rRNA gene sequences of the Food4Gut cohort (MG-RAST project ID: FOOD4GUThealthy). The remaining data supporting the findings of this study are available within the article and its supplementary materials.

## Funding

The research leading to these results was funded by the Fonds Wetenschappelijk Onderzoek – Vlaanderen (FWO) and the Fonds de la Recherche Scientifique (FNRS) under EOS Project No. 40007505, HOMISTASIS Project as well as by the Walloon Region in the context of the funding of the strategic axis FRFS-WELBIO (40009849). LBB was a Collen-Francqui Research Professor during this period of research and grateful for the support of the Francqui Fondation. EP is a PhD Research Fellow from the F.R.S.-FNRS. CDB is supported in part by the Southampton National Institute for Health and Care Research, Biomedical Research Centre (NIHR203319). PDC is honorary research director at FRS-FNRS (Fonds de la Recherche Scientifique) and the recipient of grants from FNRS (Projet de Recherche PDR-convention: FNRS T.0032.25, WELBIO-CR-2025: 40038842, EOS: HOMISTASIS program no. 40007505). MR is funded by the European Research Council (ERC) under the European Union’s Horizon 2020 research and innovation program (STOPWASTE no. 949017) and the German Research Foundation (DFG FOR5795-1_HyperMet). TC received funding from the German Research Foundation (DFG), project ID 445552570 and 564414043.

## Acknowledgments

We greatly thank Stéphanie Delieux and Bouazza Es Saadi for their precious and skilled technical assistance throughout the entire course of this work. We are grateful to the teams that shared bacterial isolates with us: Dr. Trevor D. Lawley, Dr. Hilary P. Browne and Nicholas Dawson (Wellcome Sanger Institute, UK); Dr. Petra Louis and Freda Farquharson; Dr. Mahesh Desai. We also greatly thank Dr. Nicolas Joudiou, Lionel Mignion and the UCLouvain-LDRI Nuclear and Electron Spin Technologies (NEST) platform for providing easy access and skilled assistance for the nuclear magnetic resonance spectrometry. We are also grateful to all the people that shared with us their expertise, skilled technical assistance and/or provided access to dedicated instruments.

## Author Contributions

Conception and design of the work: LBB. Design of experiments: SL and LBB. Data collection: SL, EP, AMN, SAP, AJ. Bacterial whole-genome analysis: SL with the help of TCAH, TC. Setup of the IL-6 model: PM, MR. Microbiome analysis: LBB with the support of RH. Transcriptomics analyses: AL. Sample and data collection INSYTE cohort: ES, JB, CDB. Data analysis and interpretation: SL, LBB. Participation to critical scientific discussions: AMN, PM, PDC, MR, LV, TCAH, TC, NMD. Acquisition of funding: LBB. Drafting the article: SL with the help of LBB. Critical revision of the article: all. Final approval of the version to be published: all.

## Disclosure of Interest

CDB has received research funding from Echosens, outside of this work. PDC is inventor on patent applications dealing with the use of gut bacteria and their components in the treatment of diseases. PDC was a co-founder of Enterosys. The authors declare that the research was conducted in the absence of any commercial or financial relationships that could be construed as a potential conflict of interest.

## Supplemental Figure legends

**Supplemental Figure 1. Setting the timing for the collection of the bacterial supernatants at 48h from inoculation to achieve the maximal concentration of metabolites.**

(A) Growth curve over 82 hours of four inocula of *Lachnospira eligens* C15-B4^T^ Optical density was measured at 600 nm at the timepoints indicated by a vertical dotted line. Data are presented as mean.

(B) PCA on metabolomic profile over time. ^1^H-NMR metabolomics analysis of culture medium inoculated or not with *L. eligens* C15-B4^T^ from bottles 1 and 2 presented in Figure S1A and two corresponding non inoculated bottles.

Abbreviation : *LE* (*Lachnospira eligens*).

**Supplemental Figure 2. Analysis of the whole-genome sequence of the six *Lachnospira eligens* isolates.**

(A) Results of the identification of putative antimicrobial resistance (AMR) genes, disinfectant resistance genes and plasmids present in the whole-genome sequence of the 6 *L. eligens* isolates. Detailed information about the tools and database names, versions and default parameters as well as the date of analysis and the obtained results can be found in the Supplemental Table 4.

(B) Genome comparison of the 6 *L. eligens* isolates by performing the overall genome relatedness index (OGRI) digital DNA-DNA hybridization (dDDH).

(C) Unrooted maximum likelihood phylogenetic tree of the 6 *L. eligens* isolates, inferred using Up-to-date Bacterial Core Genes (UBCGs) (concatenated alignment of 92 core genes) through the UBCG pipeline which is based on the multigene-based phylogenies methods (MBPs). Gene support indices (GSIs) and percentage bootstrap values are presented at branching points in pink and black respectively. Bar, 0.01 substitution per position.

(D) Species Genome Bin (SGB) abundance in treatment-naive AML patients (AML in orange) and healthy individuals (CT in grey). The type strain of *L. eligens* was found to belong to the SGB 5083 and the other 5 *L. eligens* isolates to the SGB 5082. Data are presented as median ± IQR. Mann-Whitney U tests, *p<0.05.

**Supplemental Figure 3. *Lachnospira eligens* CLA-AA-H260 supernatant does not reverse dexamethasone-induced myotube atrophy.**

Comparison of myotube diameter between myotubes incubated 48 hours with or without dexamethasone followed by an incubation of 48 hours (photos collected at 24 and 48 hours) with or without 1/10 diluted *L. eligens* CLA-AA-H260 supernatant filtered at 1 kDa. Data are presented as mean ± SD, representative of 1 independent experiment performed in triplicates (N = 1, n = 3). Statistical analysis of the photos collected after 24 hours of incubation : One-Way ANOVA (p=0.0019) followed by Dunnett’s multiple comparisons test, **p<0.01. Statistical analysis of the photos collected after 48 hours of incubation : One-Way ANOVA (p<0.0001) followed by Dunnett’s multiple comparisons test, ****p<0.0001.

Abbreviation : *LE* (*Lachnospira eligens*).

**Supplemental Figure 4. Superimposed ^1^H-NMR spectra of regions showing peaks of acetate, formate, lactate and ethanol.**

Superimposed ^1^H-NMR spectra were obtained from non-conditioned medium (NCM) and culture supernatant of *Lachnospira eligens (LE)* CLA-AA-H260 and *Escherichia coli* HS after they had been run or not through acid-binding columns. Spectra from Supplemental Figure 4 were analyzed to obtain the results of Figure 6A.

(A) Color legend of the superimposed ^1^H-NMR spectra.

(B) Acetate singlet at 1.93 ppm.

(C) Lactate doublet centered at 1.32 ppm.

(D) Formate singlet at 8.46 ppm.

(E) Ethanol triplet centered at 1.19 ppm.

**Supplemental Figure 5. *Serpina3n* gene expression in myotubes treated with dexamethasone with or without *Lachnospira eligens (LE)* CLA-AA-H260 supernatant.**

(A) *Serpina3n* gene expression in C_2_C_12_ myotubes (qPCR results). Data are presented as mean ± SD, representative of 1 independent experiment (N = 1, n = 3). One-Way ANOVA (p<0.0001) followed by Dunnett’s multiple comparisons test, **p<0.01, ****p<0.0001.

(B) *Serpina3n* gene expression in C_2_C_12_ myotubes (qPCR results). Data are presented as mean ± SD, representative of 1 independent experiment (N = 1, n = 3). One-Way ANOVA (p<0.0001) followed by Dunnett’s multiple comparisons test, ***p<0.001, ****p<0.0001.

**Supplemental Figure 6. Unravelling the mechanism of action from the host side – net metabolic flux analysis.**

(A) Comparison of myotube diameter between myotubes incubated with or without 1 µM of dexamethasone and with or without 1/10 diluted *Lachnospira eligens* CLA-AA-H260 supernatant. Data are presented as mean ± SD, representative of 1 independent experiment performed in triplicates (N = 1, n = 3). One-Way ANOVA (p<0.0020) followed by Dunnett’s multiple comparisons test, **p<0.01.

(B) Net production (+) or consumption (-) of acetate, formate, lactate and glucose by C_2_C_12_ myotubes following 48 hours of incubation with or without 1 µM of dexamethasone and with or without 1/10 diluted *Lachnospira eligens* CLA-AA-H260 supernatant.

Abbreviation : *LE* (*Lachnospira eligens*).

**Supplemental Figure 7. Acetate, formate, lactate and ethanol concentrations in the treatments that were orally administered to leukemic mice.**

(A) Concentration (mM) of acetate, formate, lactate and ethanol in the treatments that were orally administered to the mice. Concentrations were measured by ^1^H-NMR metabolomics. The administered non-conditioned media (NCM) and bacterial culture supernatants were concentrated using a CentriVap centrifugal concentrator. During this process ethanol evaporated.

**Supplemental Figure 8. Supplementation with *Escherichia coli* HS anoxic supernatant does not prevent liver and spleen increase in leukemic mice.**

(A) Evolution of body weight in percentage of initial body weight. Data are presented as mean ± SEM. Two-way Repeated Measures ANOVA with the Geisser-Greenhouse correction was performed followed by Tukey’s multiple comparisons test, p=ns are not shown, only the pairwise comparison of the group CT_NMC_AN vs BaF_Ecoli_AN was significant (*p<0.05).

(B) Spleen weight in percentage of initial body weight. Data are presented as mean ± SEM. One-Way ANOVA (p<0.0001) was performed followed by Tukey’s multiple comparisons test, *p<0.05, ****p<0.0001.

(D) Liver weight in percentage of initial body weight. Data are presented as median ± IQR. Kruskal-Wallis test (p<0.0001) was performed followed by Tukey’s multiple comparisons test, **p<0.01, ***p<0.001, ****p<0.0001.

**Supplemental Figure 9. Supplementation with *Escherichia coli* HS anoxic supernatant does not prevent the dysregulation of muscle atrophy markers and intestinal barrier integrity markers in leukemic mice.**

(A) Gastrocnemius muscle gene expression. Data for *Serpina3n, Trim63, Ctsl* and *Map1lc3b* are presented as median + IQR, Kruskal-Wallis test was performed (p<0.0001) followed by Dunn’s post-test. Data for *Fbxo32* are presented as mean + SEM, Welch’s ANOVA test was performed (p<0.0001) followed by Tamhane T2’s post-test. Post-tests *p<0.05, **p<0.01, ***p<0.001 and ****p<0.0001.

(B) Soleus muscle gene expression. Data for *Serpina3n* are presented as median + IQR, Kruskal-Wallis test was performed (p<0.0001) followed by Dunn’s post-test. Data for *Fbxo32* are presented as median + IQR, Kruskal-Wallis test was performed (p=0.0035) followed by Dunn’s post-test. Data for *Trim63 and Ctsl* are presented as mean + SEM, Welch’s ANOVA test was performed (p<0.0001) followed by Tamhane T2’s post-test. Data for *Map1lc3b* are presented as mean + SEM, One-way ANOVA was performed (p<0.0001) followed by Tukey’s post-test. Post-tests *p<0.05, **p<0.01, ***p<0.001 and ****p<0.0001.

(C) Jejunum gene expression. Data for *Ocln* are presented as median + IQR, Kruskal-Wallis test was performed (p=0.0012) followed by Dunn’s post-test. Data for *Tjp1* are presented as median + IQR, Kruskal-Wallis test was performed (p=0.0007) followed by Dunn’s post-test. Data for *Muc2* are presented as median + IQR, Kruskal-Wallis test was performed (p=ns). Data for *Il1b* are presented as mean + SEM, One-way ANOVA was performed (p<0.0001) followed by Tukey’s post-test. Post-tests *p<0.05, **p<0.01, ***p<0.001 and ****p<0.0001.

(D) Ileum gene expression. Data for *Ocln* and *Tjp1* are presented as mean + SEM, One-way ANOVA was performed (p=ns). Data for *Fbxo32* are presented as median + IQR, Kruskal-Wallis test was performed (p=0.0035) followed by Dunn’s post-test. Data for *Il1b* are presented as mean + SEM, One-way ANOVA was performed (p=0.0092) followed by Tukey’s post-test. Post-tests *p<0.05, **p<0.01, ***p<0.001 and ****p<0.0001.

Abbreviations *: Fbxo32* (F-box protein 32 (Atrogin-1)), *Trim63* (Tripartite motif containing 63 (Murf-1)), *Ctsl* (Cathepsin L), *Map1lc3b* (Microtubule associated protein 1 light chain 3 beta), *Serpina3n* (Serpin family A member 3N), *Muc2* (Mucin 2), *Ocln* (Occludin), *Tjp1* (Tight junction protein 1 (ZO-1)), *Il1b* (Interleukin 1 beta).

**Supplemental Figure 10. Acetate, formate and lactate levels in the systemic blood of mice at necropsy.**

Area Under the Curve (AUC) of acetate, formate and lactate in the systemic blood of mice at the day of necropsy. Concentrations were measured by ^1^H-NMR metabolomics. Data are presented as mean ± SEM. One-Way ANOVA was performed and was not significant for acetate and formate. For lactate, One-Way ANOVA (p=0.0001) was followed by Tukey’s multiple comparisons test, **p<0.01, ****p<0.0001.

## Supplemental Table legends

**Supplemental Table 1. Information about *Lachnospira eligens* isolates.**

**Supplemental Table 2. Primer sequences.**

**Supplemental Table 3. *CheckM* output (v1.2.2 ; lineage specific workflow).**

**Supplemental Table 4. *CheckM2* output (v1.0.2).**

**Supplemental Table 5. Antimicrobial Resistance Genes (AMR) identification in *Lachnospira eligens* isolates.**

