## Supplemental Material and Methods for "Mixed acid fermentation products from *Lachnospira eligens* counteract myotube atrophy"

### Supplemental Materials and Methods

#### Gut microbiota analysis

***Data collection – MicroAML, Wang, Food4Gut and INSYTE (INvestigation of SYnbiotic TreatmEnt in metabolic dysfunction-associated steatotic liver disease (MASLD)) cohorts***

Metagenomic sequencing data and/or 16S rRNA gene sequencing data were collected from different cohorts, namely MicroAML cohort [1] , the Wang cohort [2], the Food4Gut cohort [3] and the INSYTE cohort [4, 5] were described in the original articles. The MicroAML study is a Belgian multi-centric, prospective, observational, clinical study (ClinicalTrials.gov Identifier NCT03881826) which aimed to assess the composition and activity of the gut microbiota of 30 treatment-naïve patients with newly acute myeloid leukemia (AML) compared to 30 (1:1) healthy paired individuals. The Wang cohort is part of a Chinese study which aimed to assess the relationship between AML and intestinal barrier function, and more specifically the role of butyrate, in 31 treatment-naïve patients with AML compared to 30 healthy individuals. The Food4Gut study is a Belgian single group-design trial (ClinicalTrials.gov Identifier NCT03540550) which aimed to assess the impact of consuming inulin-type fructans-rich vegetables daily on gut microbiota, gastro-intestinal symptoms, and food-related behavior in 26 healthy individuals. The INSYTE study is an English randomized double-blind placebo-controlled trial (ClinicalTrials.gov Identifier NCT01680640) in 104 patients with MASLD which aimed to evaluate the effect of a 12-14 months synbiotic intervention (synbiotic n = 55 or placebo n = 49) on liver fat, liver fibrosis and gut microbiota. In the current work, only the data collected at baseline were used for the Food4Gut study and the INSYTE study.

***16S rRNA gene amplicon sequencing – bioinformatics – MicroAML, Wang and Food4Gut cohorts***

The same bioinformatics pipeline was applied to the data collected from the three cohorts to allow comparisons between cohorts. Bioinformatics analyses were performed *in-house* as previously described [1]. Initial quality filtering of the reads was performed with the Illumina Software, yielding an average of 84 900 reads (SD 20 869 reads) per sample for the MicroAML cohort, an average of 11 1507 reads (SD 19 690 reads) per sample for the Food4Gut cohort and an average of 84 448 reads (SD

19 447 reads) per sample for the INSYTE cohort. 59 046 reads (SD 11 058 reads) were available for the Wang cohort. Quality scores were visualized with the *FastQC* software (http://www.bioinformatics.babraham.ac.uk/ publications.html), and reads were trimmed to 220 bp (R1) and 200 bp (R2) with the *FASTX-Toolkit* (http://hannonlab.cshl.edu/fastx_toolkit/). Next, reads were merged with the merge-illumina-pairs application v1.4.2 (with P = 0.03, enforced Q30 check, perfect matching to primers which are removed by the software, and otherwise default settings including no ambiguous nucleotides allowed) [6]. The *UPARSE* pipeline implemented in *USEARCH* v11 [7]) was used to further process the sequences. Amplicon sequencing variants (ASVs) were identified using *UNOISE3* [8]). The analysis allowed the determination of 4 134 ASVs (MicroAML cohort), 1 733 ASVs (Wang cohort), 2 049 ASVs (Food4Gut cohort) and 2 717 ASVs (INSYTE cohort). Two to three ASVs were identified as *Lachnospira eligens* in each of these datasets using *blast* (version 2.9.0). Criteria for identification were set at a minimum of 99% identity with 0 to 1 mismatch and only ASV with a similarity score of 1 with *L. eligens* on RDP Seqmatch were considered as assigned to *L. eligens*. The relative abundances of ASV assigned to *L. eligens* were summed up to determine *L. eligens* relative abundance. For the INSYTE and Food4Gut cohorts, only samples collected at baseline before nutritional interventions were considered.

##### *Metagenomic sequencing – bioinformatics – MicroAML cohort*

A new set of analyses using updated tools and databases was performed on the previously published metagenomics data [1] using the UCLouvain SSS high-performance cluster. An average of 67 million paired reads per sample (SD 8 million) were generated. *Trimmomatic* (version 0.39) [9]) was used to trim adapters and low-quality reads (average quality scores < 20, SLIDINGWINDOW:6:20) and only reads with a minimum length of 100 bp were kept for the downstream analysis. *Kneaddata* (version 0.10.0) (https://huttenhower.sph.harvard.edu/kneaddata) was used to trim repetitive sequences and filter out human DNA reads based on the human reference genome (hg37dec_v0.1) thanks to an implementation of *Bowtie2* [10]. *MetaPhlAn4* (version 4.1.1) was used to estimate the taxonomic composition of the gut microbiome using the *CHOCOPhlAnSGB* database (version vJun23_202403). The SGB profiles were transformed into GTDB profiles using the *sgb_to_gtdb_profile.py* function in *MetaPhlAn4*.

##### *Metagenomic sequencing – exploration of taxonomical data*

For each sample, the cumulative relative abundance of taxa that were associated with different oxygen tolerances was determined. The level of facultative anaerobes, aerobes, obligate aerobes, microaerophiles, anaerobes and obligate anaerobes were computed based on an aggregation at the species level and the oxygen sensitivity retrieved from *BacDive* database on June 12, 2025 [11].

The cumulative relative abundance of oral species was computed by summing up the relative abundance of species whose primary body site is the mouth, as registered in the expanded *Human oral microbiome database* V4 (*eHOMD*) [12]. Among the 1942 species present in MicroAML samples, 89 were indicated as species for which primary body site is the mouth.

#### $^{1}$H-NMR metabolomics analyses

##### *Sample preparation*

Bacterial supernatant and NMC samples were prepared as follows: 400 µL of sample was mixed with 200 µL of NMR buffer ($D_2O$, pH = 7 ($KHPO_4$-$K_2HPO_4$ 1.5 M), trimethylsilylpropionic acid Na Salt D4 99.80%D (TSP) (Eurisotop, France) 2 mM as standard). The mix was then transferred into 5 mm diameter NMR tubes.

Blood from Balb/c mice was prepared as follows: 40 µL of sample was mixed with 20 µL of NMR buffer ($H_2O$-$D_2O$ (1:1), pH = 7 ($NaH_2PO_4$- $Na_2HPO_4$ 0.2 M), trimethylsilylpropionic acid Na Salt D4 99.80%D (TSP) (Eurisotop, France) 2 mM as standard). The mixture was then centrifuged (5 minutes, 12 000g, 4°C) and transferred into 1.7-mm-diameter NMR tubes.

##### *Data collection*

NMR data were acquired on a Bruker Avance 600 MHz NMR spectrometer equipped with a cryoprobe. During acquisition, sample temperature was maintained at 300 K. Spectra were collected with a 1D NOESY pulse sequence for the supernatant and NCM samples and with a 1D CPMG pulse sequence for blood samples. The 1D NOESY pulse sequence covered 21 ppm. Spectra were digitized in 65K data points during a 2.6 s acquisition time. The mixing time was set to 10 ms, and the relaxation delay between scans was set to 4 s. The 1D CPMG pulse sequence covered 20 ppm. Spectra were digitized in 65K data points during a 2.7 s acquisition time. The relaxation delay between scans was set to 4 s.

Spectra were acquired using 128 scans for the supernatant and NCM samples and 256 scans for blood samples.

##### *Data processing and analysis*

The data were processed using *MestReNova* (version 14.3.0). The spectra were zero filled with a factor of two. They were submitted to apodization using a 0.3 Hz decaying exponential function and fast Fourier transformed. Automated phase correction and second-order polynomial baseline correction were applied to all samples.

The spectra obtained from bacterial supernatant and NMC samples were aligned on TSP, which was used as a chemical shift and quantification reference for all spectra. *Chenomx NMR Suite* (version 8.6) was used to identify and quantify metabolites. Quantitative fitting of each spectrum was carried out in batch mode, followed by manual adjustment.

To obtain the Principal Component Analysis (PCA) on metabolic profile over time of the supernatant of *L. eligens* C15-B4$^{T}$ and NCM, only the region from 0.13 to 10 ppm was conserved and the water signal was removed. Optimized bucketing [13] was performed using *MATLAB* (version 9.2) followed by a PCA, which was performed using the function *pca* of the *mixOmics* R package [14].

The spectra obtained from Balb/c mice blood samples were processed and analyzed differently than those acquired from bacterial supernatant and NCM samples due to interactions between blood albumin and the quantification reference TSP, which caused line broadening and loss of the TSP signal. Consequently, absolute quantification of metabolites was not possible. The spectra obtained from Balb/c mice blood samples were aligned on formate and only the region from 0.4 to 8.55 ppm was conserved, the water signal was removed as well as the following regions of glucose signal 3.95-3.715 ppm and 3.58-3.4 ppm. Then constant sum normalization was performed. *Chenomx NMR Suite* (version 8.6) was used to identify and quantify area under the curve (AUC) of metabolites. The fitting of each spectrum was carried out in batch mode, followed by manual adjustment.

#### Transcriptome analyses

##### *C2C12 whole transcriptome analysis*

RNAseq analysis was performed on RNA isolated from myotubes from two independent experiments. Total RNA was isolated from C2C12 myotubes by TriPure reagent (Roche, Switzerland) and RNA quantity was evaluated using a NanoPhotometer Spectrophotometer (Implen, Germany). The quality of the RNA samples was assessed using a 2100 Bioanalyzer System (Agilent Technologies, USA). All RIN values were above 7, supporting RNA integrity. Illumina TruSeq Stranded mRNA libraries were prepared using polyA tail selection. Following library quality control, libraries were pooled and sequenced using a 2 × 150bp paired-end configuration on a NovaSeq X instrument (Macrogen, The Netherlands). FASTQ files were processed using a standard RNAseq pipeline including *Trimmomatic* (version 0.39) [9] to remove low quality reads and *HISAT2* (version 2.2.1) [15] to align reads to the mouse reference genome (GRCm38). Gene expression levels were quantified using *featureCounts* from *Subread* (version 2.0.3) [16] and genomic features were defined based on the *Ensembl Mus_musculus.GRCm38.94.gtf* annotation file. Differential expression analyses were performed with *DESeq2 Bioconductor* package (version 1.36.0) [17], using a design formula of the form ~ Group + experiment, thereby accounting for both treatment and variability between the two independent experiments. Genes with an adjusted p-value $< 0.05$ and a log2 fold change $> 1$ were considered differentially expressed.

##### *Tissue and C2C12 mRNA analysis by qPCR*

Total RNA was isolated from tissue or C2C12 myotubes by TriPure reagent (Roche, Basel, Switzerland). cDNA was prepared by reverse transcription of 1 µg total RNA using the Goscript RT Mix OligoDT kit (Promega, USA) with a Biometra TOne instrument and software (Analytik Jena, Germany). Real-time polymerase chain reactions (PCR) were performed with a StepOnePlus/QuantStudio Real-Time PCR System and software (Applied Biosystems, Den Ijssel, The Netherlands) or a CFX96 TouchTM instrument and software (Bio-Rad Laboratories, CA, USA) using the GoTaq® qPCR Master Mix (Promega, USA). To measure the expression of *Serpina3n* the iQ™ SYBR® Green Supermix (Bio-Rad, USA) was used. All samples were run in duplicate in a single 96-

well reaction plate, and data were analyzed according to the $2^{-\Delta\Delta CT}$ method. The purity of the amplified product was verified by analyzing the melt curve performed at the end of amplification. The ribosomal protein L6 (*Rpl6*) gene was used as housekeeping gene. The primer sequences for the targeted mouse genes are detailed in Supplemental Table 2.
