## Supplemental Figures for "Mixed acid fermentation products from *Lachnospira eligens* counteract myotube atrophy"

### Supplemental Figure 1

## A

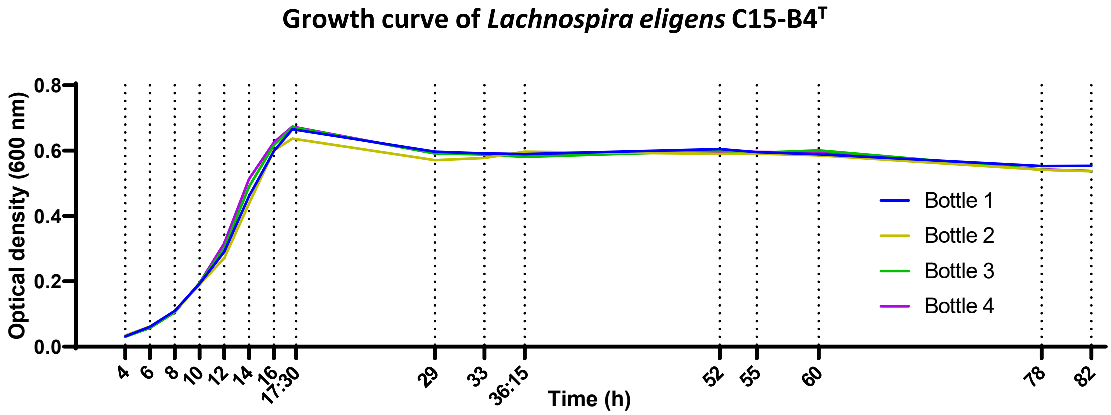


## B

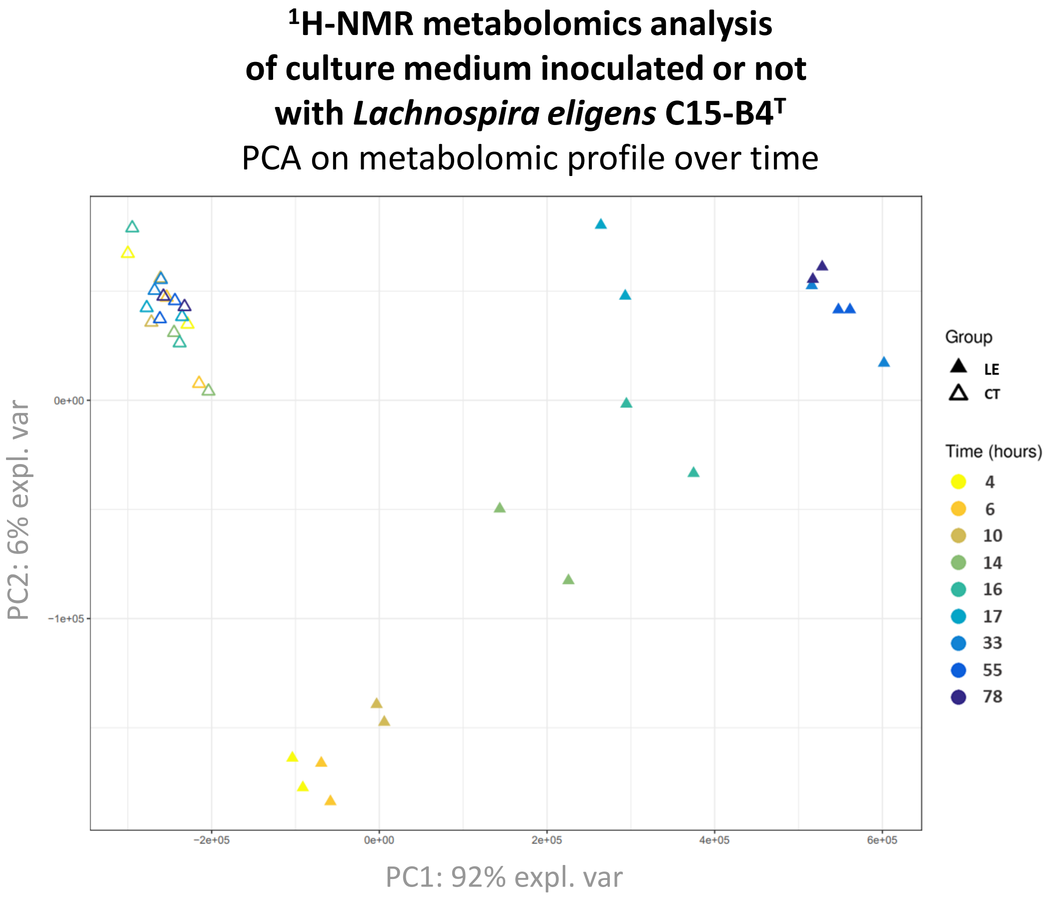

### Supplemental Figure 2

**A**

| Tool name | Database name | C15-B4^T | H1-24 | CLA_AA_H260 | I42 | WS1 | WS2 |
|---|---|---|---|---|---|---|---|
| ResFinder v.4.6.0 | ResFinder v. 2.4.0 | 0 | 0 | 0 | 0 | 0 | 0 |
| | DisinFinder v. 2.0.1 | 0 | 0 | 0 | 0 | 0 | 0 |
| RGI v.6.0.3 | CARD v.3.2.8 | 0 | 0 | 0 | 0 | 0 | 0 |
| ABRicate (in Galaxy) v.1.0.1 | ARG-ANNOT | 0 | 0 | 0 | 1 AMR gene : (Gly)vanA-G | 0 | 0 |
| | NCBI Bacterial Antimicrobial Resistance Reference Gene Database | 0 | 0 | 0 | 0 | 0 | 0 |
| | VFDB | 0 | 0 | 0 | 0 | 0 | 0 |
| | Megares 2.00 | 0 | 0 | 0 | 0 | 0 | 0 |
| PlasmidFinder v.2.0.1 | PlasmidFinder | 0 | 0 | 0 | 0 | 0 | 0 |

**B**

dDDH (digital DNA-DNA hybridization)

| Compared strains | dDDH (in %) | Confidence interval (in %) |
|---|---|---|
| H1-24 vs WS2 | 100.0 | [100.0 – 100.0] |
| CLA-AA-H260 vs WS2 | 88.5 | [86.0 – 90.6] |
| H1-24 vs CLA-AA-H260 | 88.5 | [86.0 – 90.6] |
| I42 vs CLA-AA-H260 | 88.0 | [85.5 – 90.1] |
| I42 vs WS2 | 86.3 | [83.7 – 88.6] |
| I42 vs H1-24 | 86.3 | [83.7 – 88.6] |
| CLA-AA-H260 vs WS1 | 85.9 | [83.2 – 88.2] |
| H1-24 vs WS1 | 85.8 | [83.2 – 88.1] |
| WS1 vs WS2 | 85.8 | [83.2 – 88.1] |
| I42 vs WS1 | 85.0 | [82.3 – 87.4] |
| C15-B4^T vs WS1 | 58.5 | [55.7 - 61.3] |
| C15-B4^T vs CLA-AA-H260 | 58.4 | [55.6 – 61.2] |
| C15-B4^T vs H1-24 | 58.0 | [55.2 – 60.7] |
| C15-B4^T vs WS2 | 58.0 | [55.2 – 60.7] |
| C15-B4^T vs I42 | 57.8 | [55.0 – 60.6] |

**C**

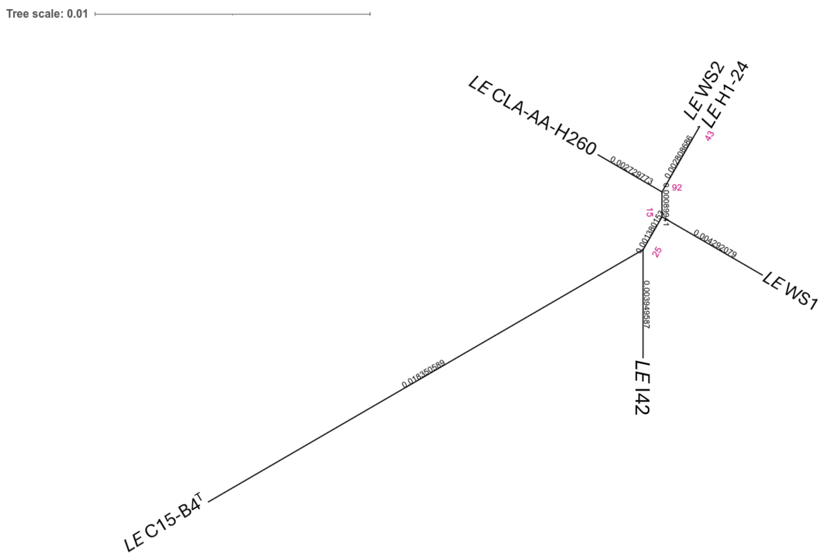


**D**

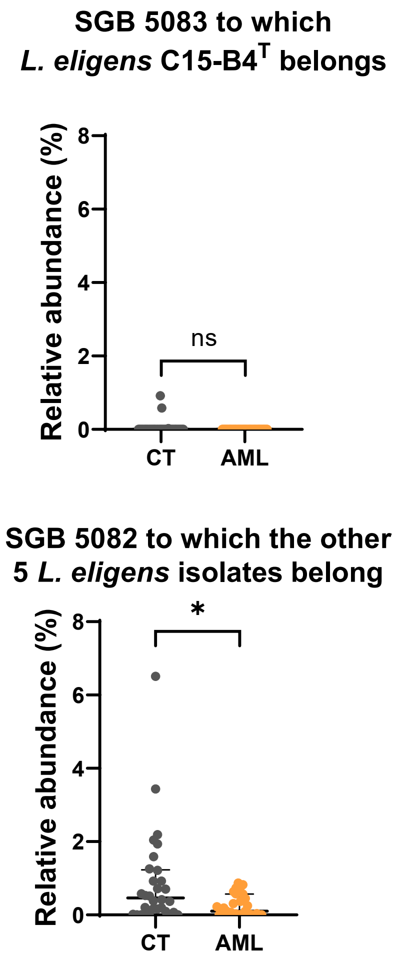

### Supplemental Figure 3

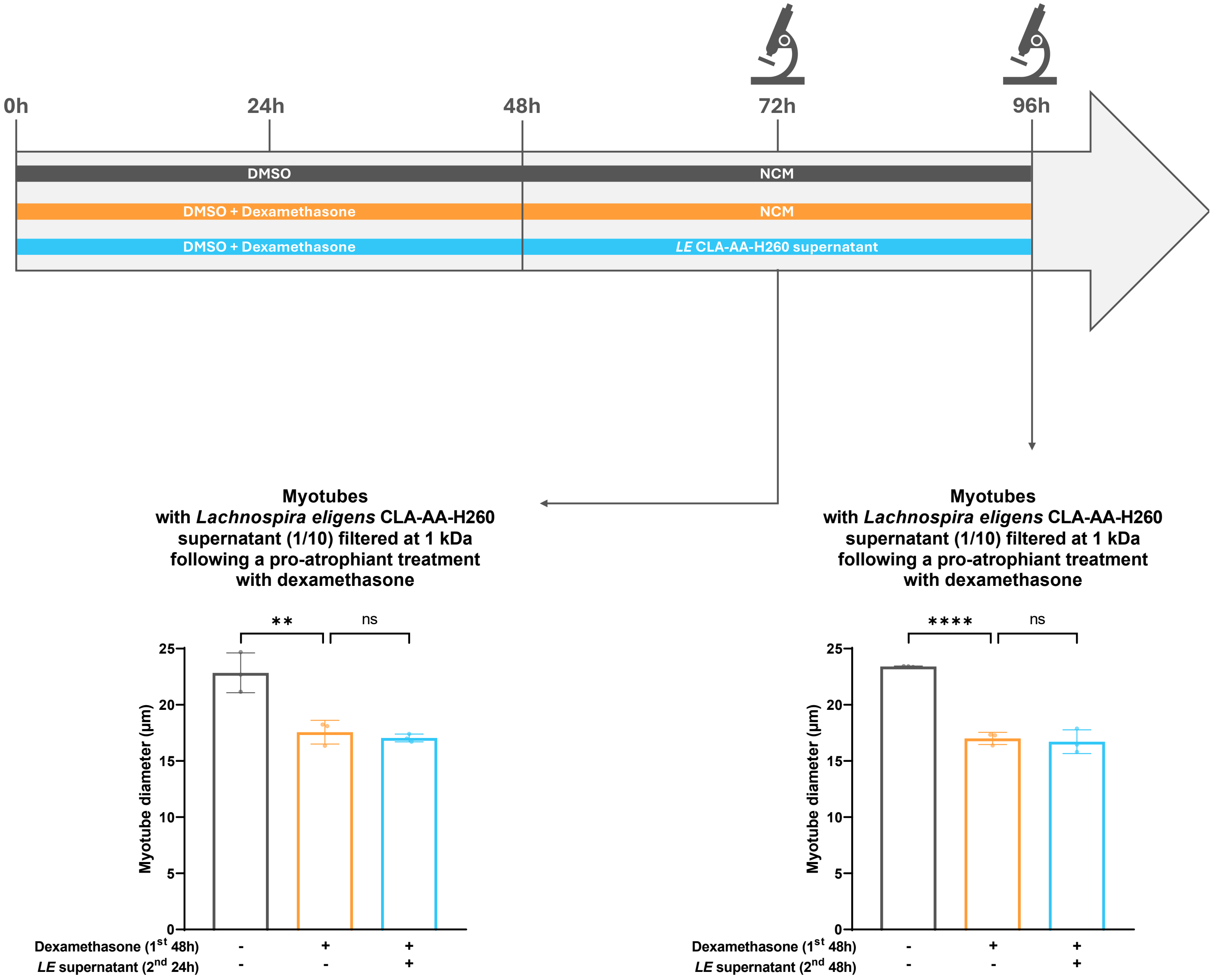

#### Supplemental Figure 4

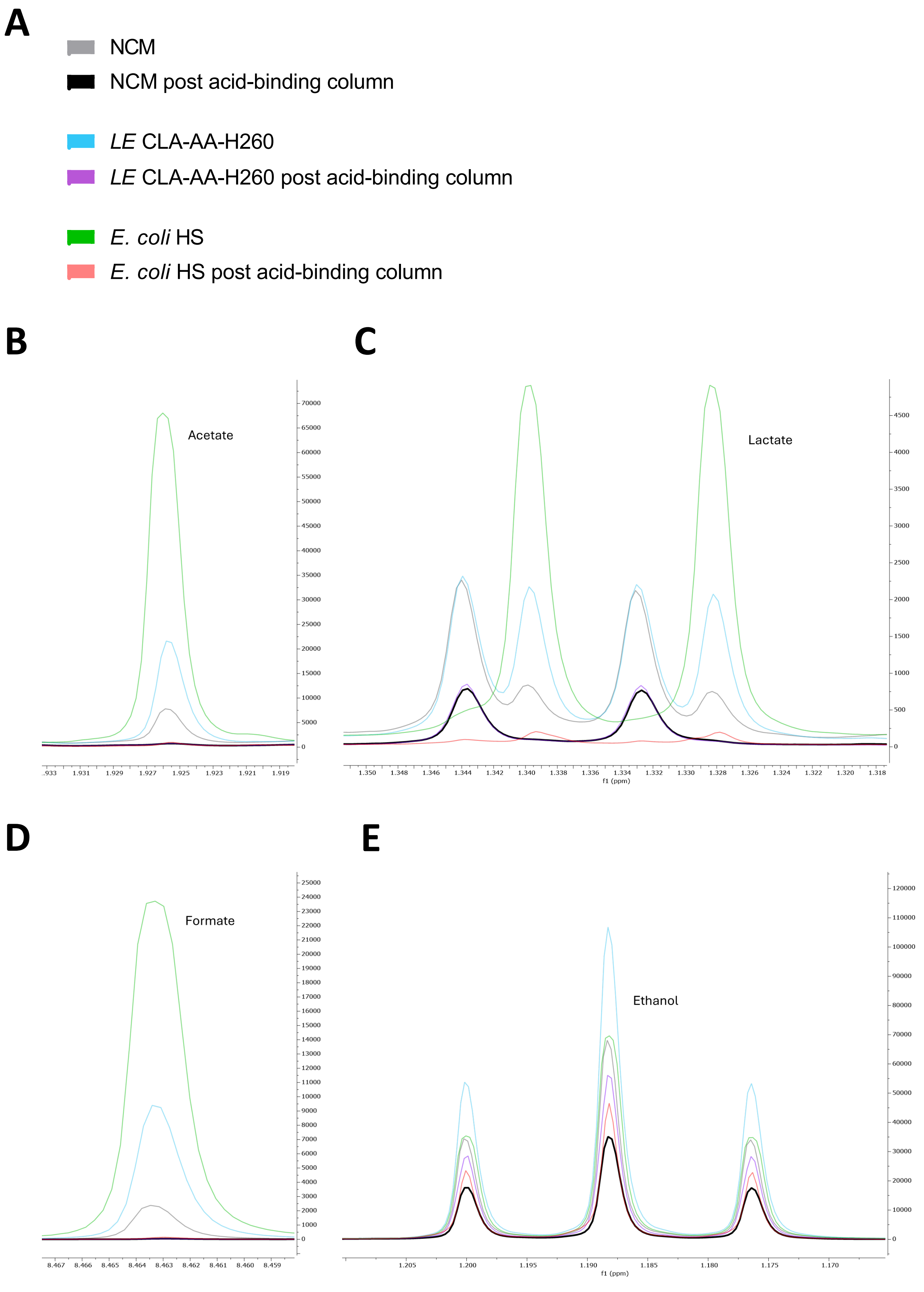

### Supplemental Figure 5

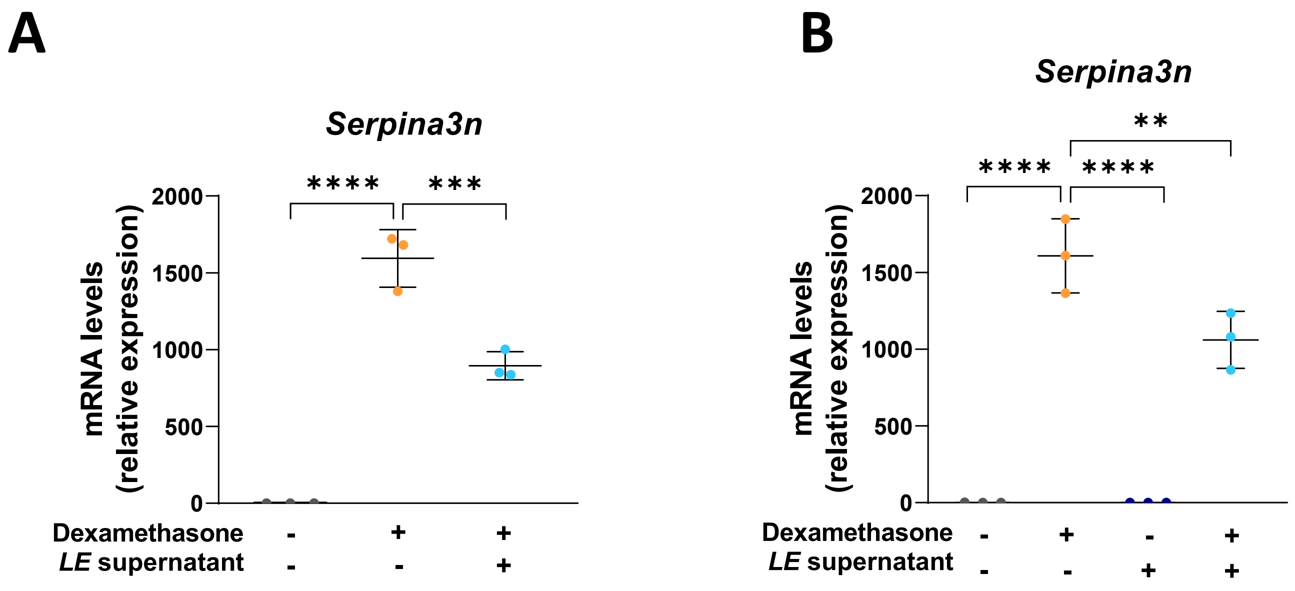

### Supplemental Figure 6

## A

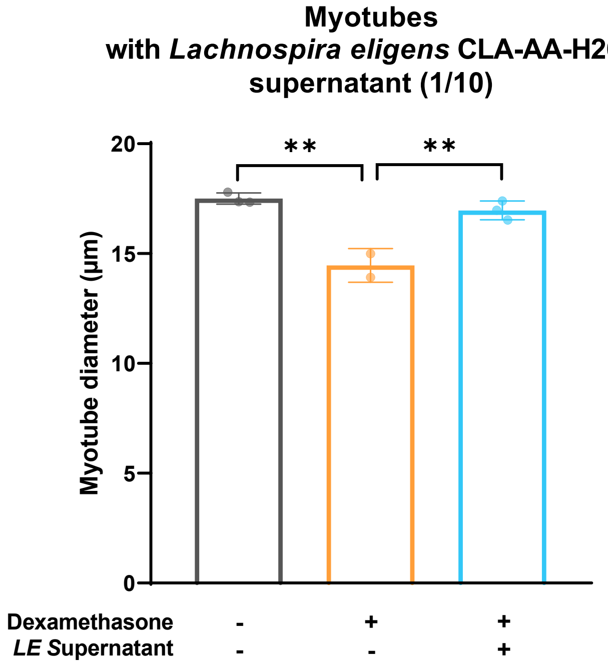


## B

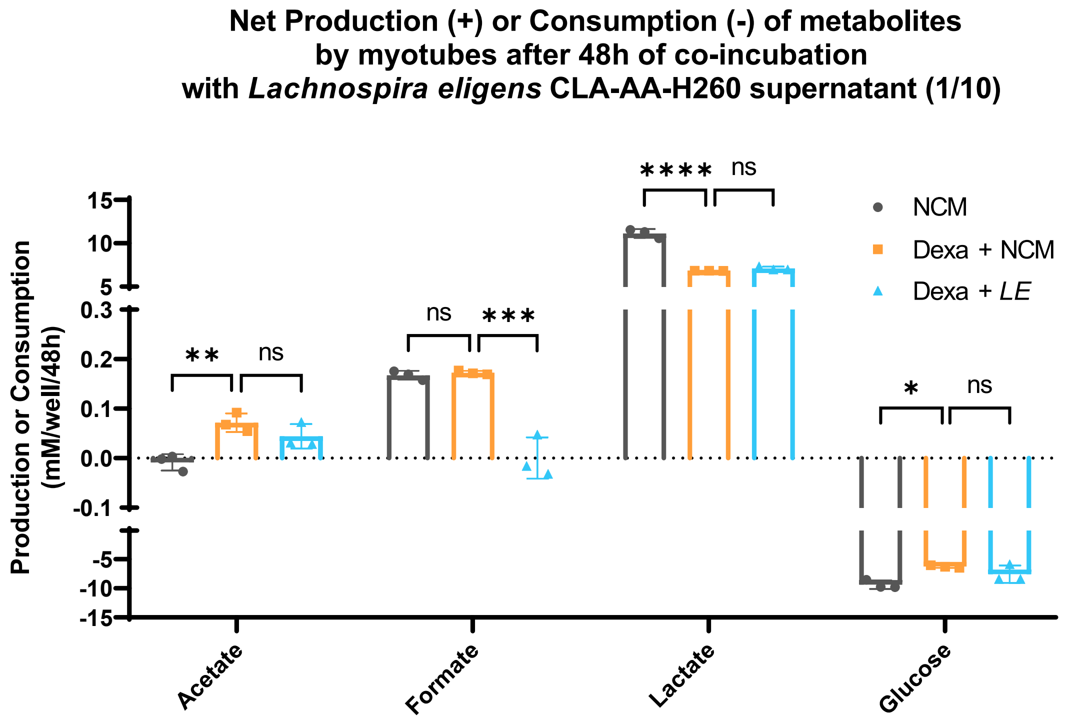

### Supplemental Figure 7

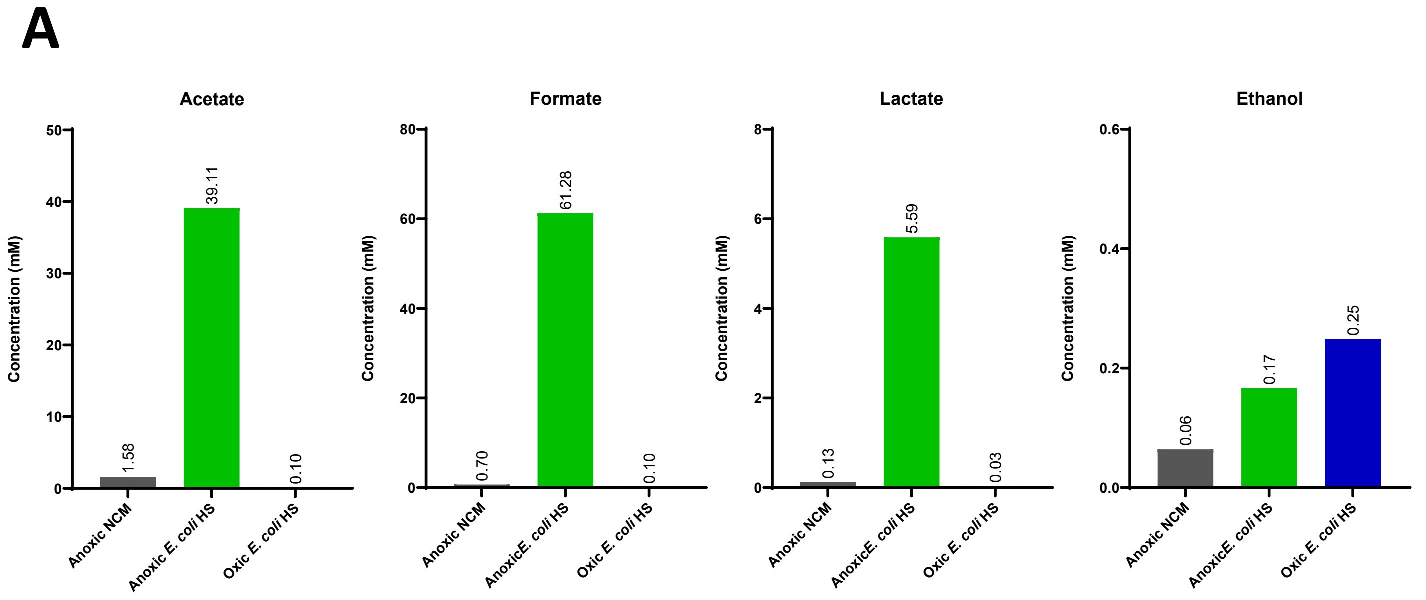

### Supplemental Figure 8

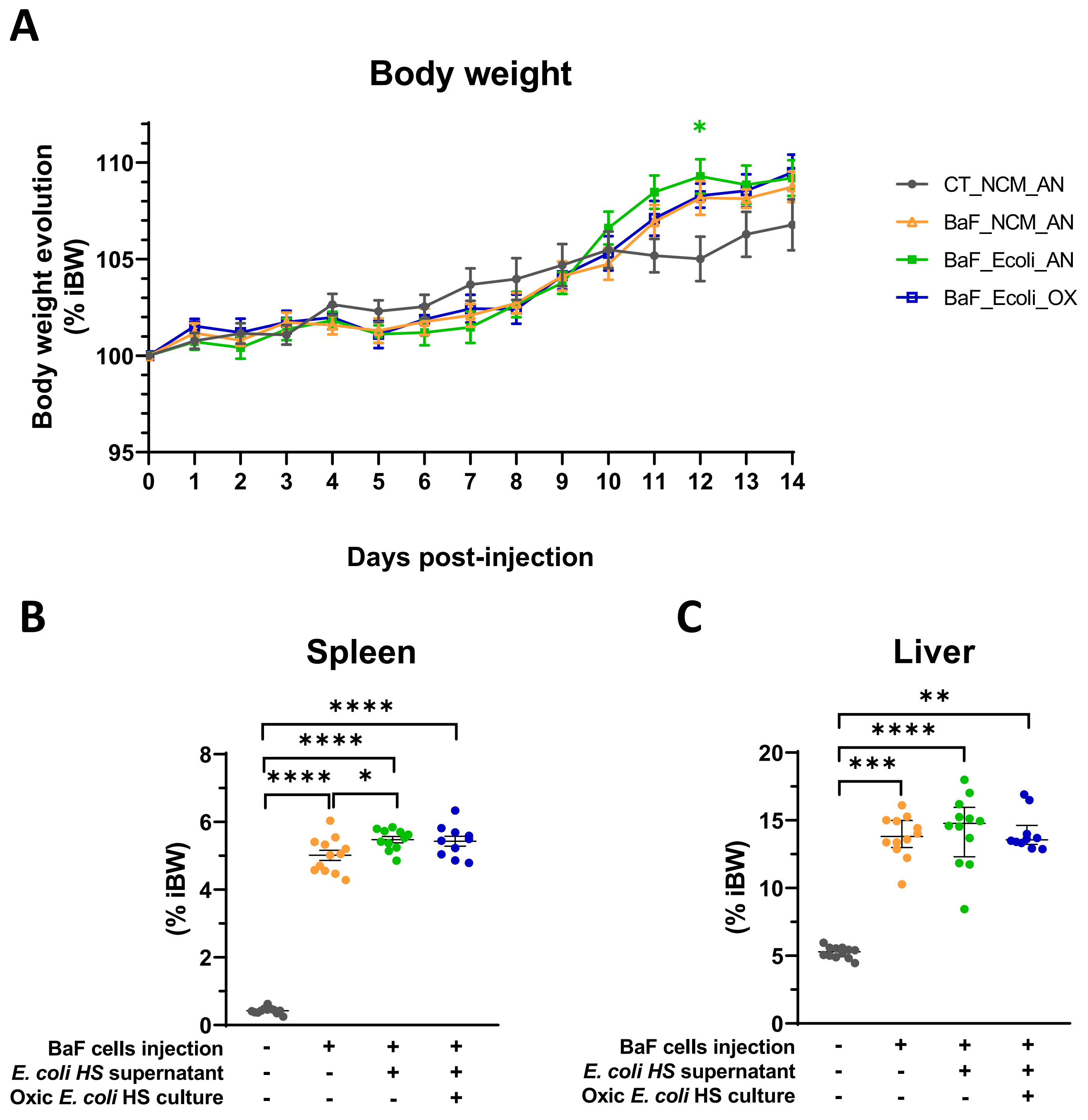

### Supplemental Figure 9

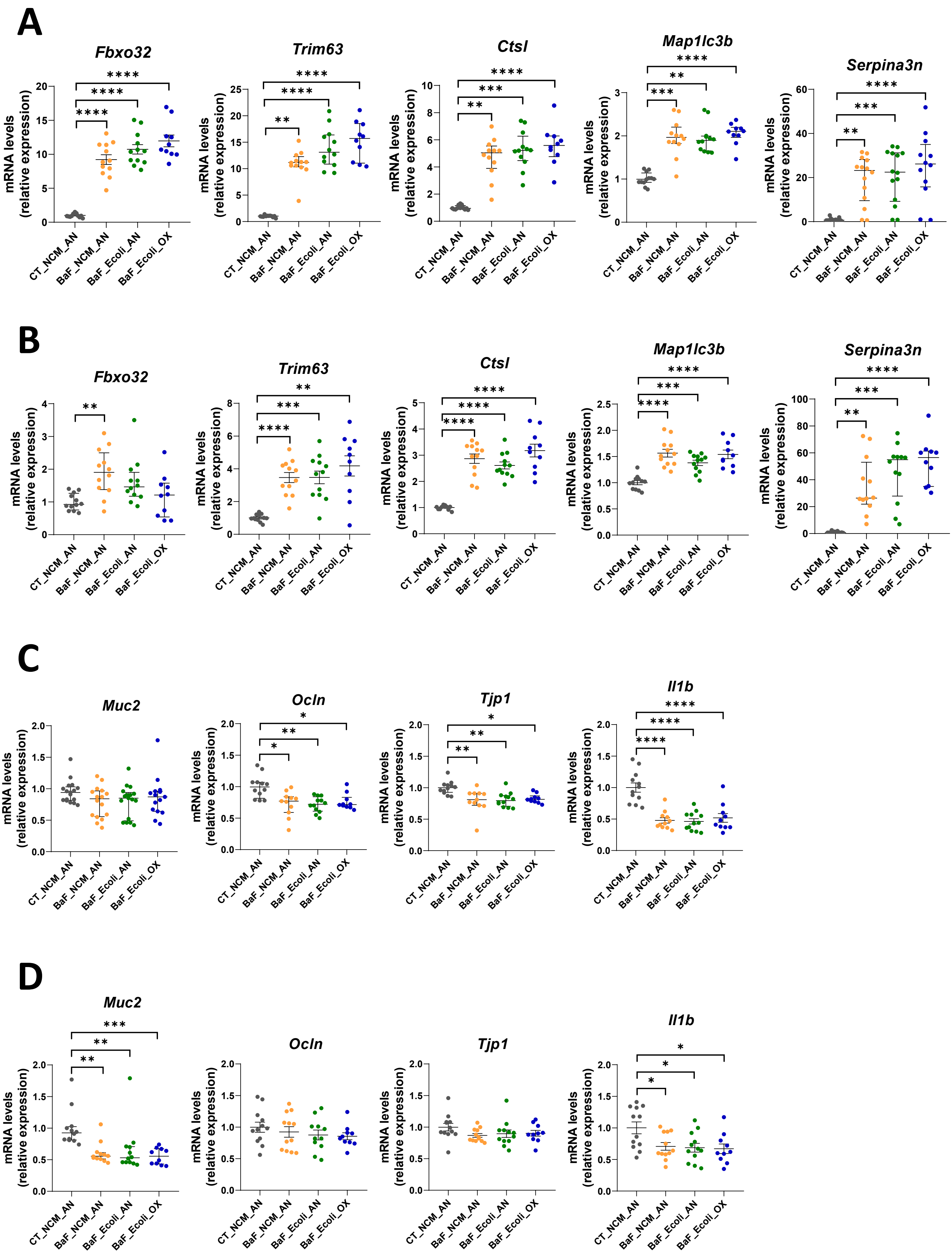

### Supplemental Figure 10

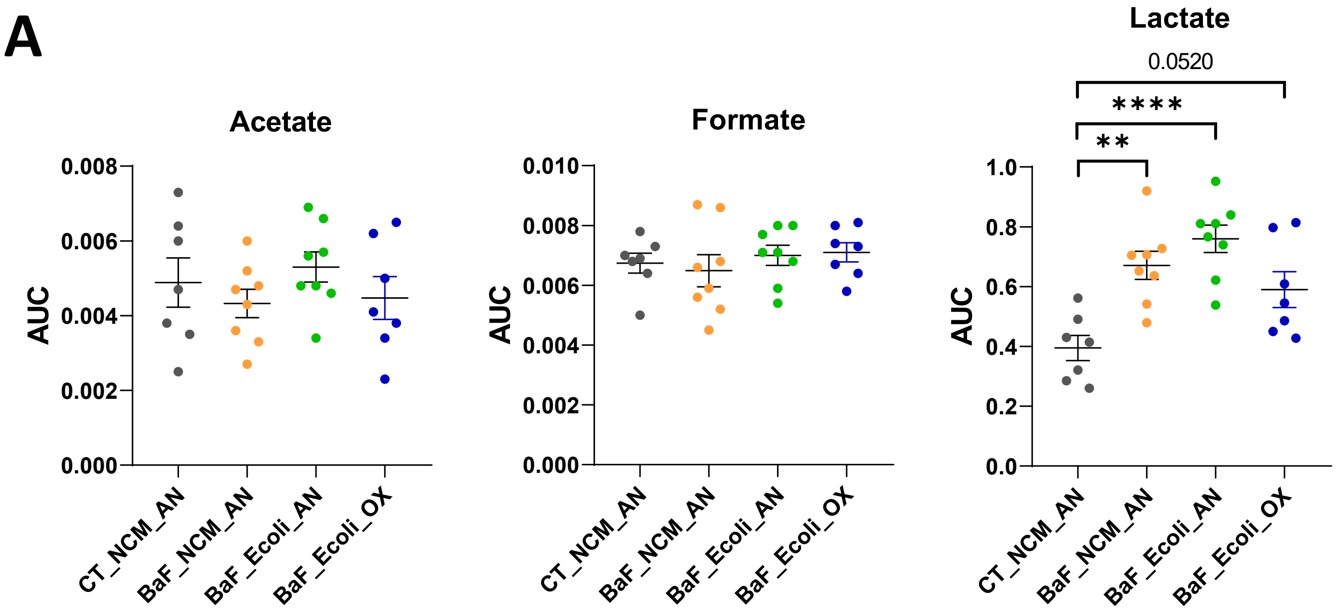
